# Selective abrogation of S6K2 maps lipid homeostasis as a survival vulnerability in MAPKi-resistant NRAS^MUT^ melanoma

**DOI:** 10.1101/2021.04.07.438684

**Authors:** Brittany Lipchick, Adam N. Guterres, Hsin-Yi Chen, Delaine M. Zundell, Segundo Del Aguila, Patricia I. Reyes-Uribe, Subhasree Basu, Xiangfan Yin, Andrew V. Kossenkov, Yiling Lu, Gordon B. Mills, Qin Liu, Aaron R. Goldman, Maureen E. Murphy, David W. Speicher, Jessie Villanueva

## Abstract

Although oncogenic NRAS activates MAPK signaling, inhibition of the MAPK pathway is not therapeutically efficacious in NRAS-mutant tumors. Here we report that silencing the ribosomal protein S6 kinase 2 (S6K2), while preserving the activity of S6K1, perturbs lipid metabolism, enhances fatty acid unsaturation, and triggers lethal lipid peroxidation selectively in NRAS-mutant melanoma cells that are resistant to MAPK inhibition. S6K2 depletion induces ER stress, and PPARα activation, triggering cell death selectively in MAPKi-resistant melanoma. We show that combining PPARα agonists and polyunsaturated fatty acids phenocopies the effects of S6K2 abrogation, blocking tumor growth in PDX and immunocompetent mouse pre-clinical models. Collectively, our study establishes S6K2 and its effector subnetwork as promising targets for NRAS-mutant melanoma that are resistant to global MAPK pathway inhibitors.

**One Sentence Summary:** S6K2 is a vulnerability in MAPK inhibitor-resistant NRAS-mutant melanoma

## INTRODUCTION

RAS-mutant tumors are highly aggressive and mostly refractory to currently available targeted therapies. In cutaneous melanoma, mutations in NRAS occur in almost 30% of all tumors(*1*). Oncogenic NRAS elicits persistent activation of the RAF/MEK/ERK (MAPK) cascade, which plays a key role in melanomagenesis; therefore, inhibition of MAPK has been evaluated as a potential therapeutic approach for NRAS-mutant melanoma. However, inhibitors of the MAPK pathway as single agents elicit less than 20% response rate and do not prolong the survival in NRAS-mutant melanoma patients compared to chemotherapy(*2, 3*). Further, suppression of MAPK signaling often leads to feedback or compensatory activation of the PI3K/AKT pathway(*4–7*). Unfortunately, drug combinations that inhibit both the MAPK and PI3K pathways are poorly tolerated in patients, and clinically efficacious doses have not been achieved(*8–12*). Although simultaneous inhibition of the MAPK and PI3K pathways is generally toxic, there is no evidence that the proximal molecular causes of that toxicity are the same in normal and in cancer tissues. This raises the possibility that targeting a convergent subnetwork or node, downstream from the PI3K and MAPK pathways, could elicit lethality in melanoma cells, while sparing normal tissues. Thus, mapping out the MAPK/PI3K downstream subnetwork could help identify much needed actionable drug targets for NRAS-mutant tumors.

Dual inhibition of MEK and PI3K in NRAS-mutant melanoma pre-clinical models leads to substantial perturbation of metabolic pathways(*13*). Oncogenic NRAS controls cell metabolism through the mechanistic target of rapamycin protein complex 1, mTORC1, a critical RAS effector(*14*) that integrates upstream signals, including the MAPK and PI3K pathways(*15, 16*). Key effectors of mTORC1 are the 40S ribosomal kinases (S6K), S6K1 and S6K2(*17*). The S6K pathway plays important roles in gene transcription, protein translation, cellular metabolism, and cell survival(*16–18*). Despite the high degree of structural homology between S6K1 and S6K2, their expression, activation and cellular localization are differentially regulated(*17*). Hyperactivated S6K2 signaling is characteristic of breast cancer subtypes(*19*), with S6K2 expression promoting breast cancer cell survival(*20*). Notably, loss of S6K1 typically leads to compensatory upregulation of S6K2 and vice versa(*21–24*). Furthermore, S6K1 and S6K2 have overlapping, as well as distinct biological functions(*16–18, 23–25*). This suggests that the functional activity of these two kinases is tightly coordinated and imbalance between these two kinases could lead to differential outcomes.

Here, we identified a S6K2/PPARα subnetwork downstream of the MAPK and PI3K pathways. Perturbation of this subnetwork by uncoupling S6K1 and S6K2 triggers a lipid metabolic imbalance, leading to oxidative cell death selectively in NRAS-mutant melanomas that are resistant to MAPK inhibition. Further, we provide proof-of-concept that this vulnerability can be exploited to curb MAPKi-resistant tumors and to inform the design of future strategies to combat NRAS-mutant melanoma.

## RESULTS

### S6K2 is a vulnerability in NRAS-mutant melanoma resistant to MAPK inhibitors

Since MAPK inhibition elicits heterogeneous effects in NRAS-mutant melanoma(*26*), we sought to identify potential vulnerabilities in MAPK inhibitor (MAPKi)-resistant tumor cells. We first analyzed the effect of MEK (trametinib and MEK162) or ERK (SCH-984 and BVD-523) inhibitors, collectively referred to as MAPKi, on viability of NRAS-mutant melanoma cells (Fig. S1A-C, Supplementary Table 1). MAPKi induced variable effects on viability, proliferation, and cell death (Fig. S1A-E), despite effective and persistent inhibition of the MAPK pathway (Fig. S1F). We classified cells as MAPKi-sensitive (MAPKi-S) based on their response to MAPKi, defined as ≥75% suppression of BrdU incorporation and ≥ 35% cell death (Fig. S1D-E). Conversely, MAPKi elicited modest or no suppression of BrdU (<75%; Fig. S1D) and marginal induction of cell death (< 35%; Fig. S1E) in the MAPKi-resistant (MAPKi-R) cells.

To further explore signaling perturbations in MAPKi-R *vs.* MAPKi-S cells, we analyzed cells treated with either the MEK inhibitor trametinib (Tram) or the ERK inhibitor SCH772984 (SCH984) by Western blotting and RPPA (Fig. 1A, Fig. S1F-G and Supplementary Table 2). MAPKi similarly suppressed the MAPK pathway in both sensitive and resistant cells. MAPKi-treatment did not induce significant differential activation of PI3K/AKT (Fig. 1A, Supplementary Table 2). In contrast, MAPKi treatment blocked phosphorylation of S6 kinase (S6K) and its substrate S6 in MAPKi-S, but not in MAPKi-R melanoma cells (Fig. 1A, Fig. S1F and Supplementary Table 2).

**Fig. 1.**
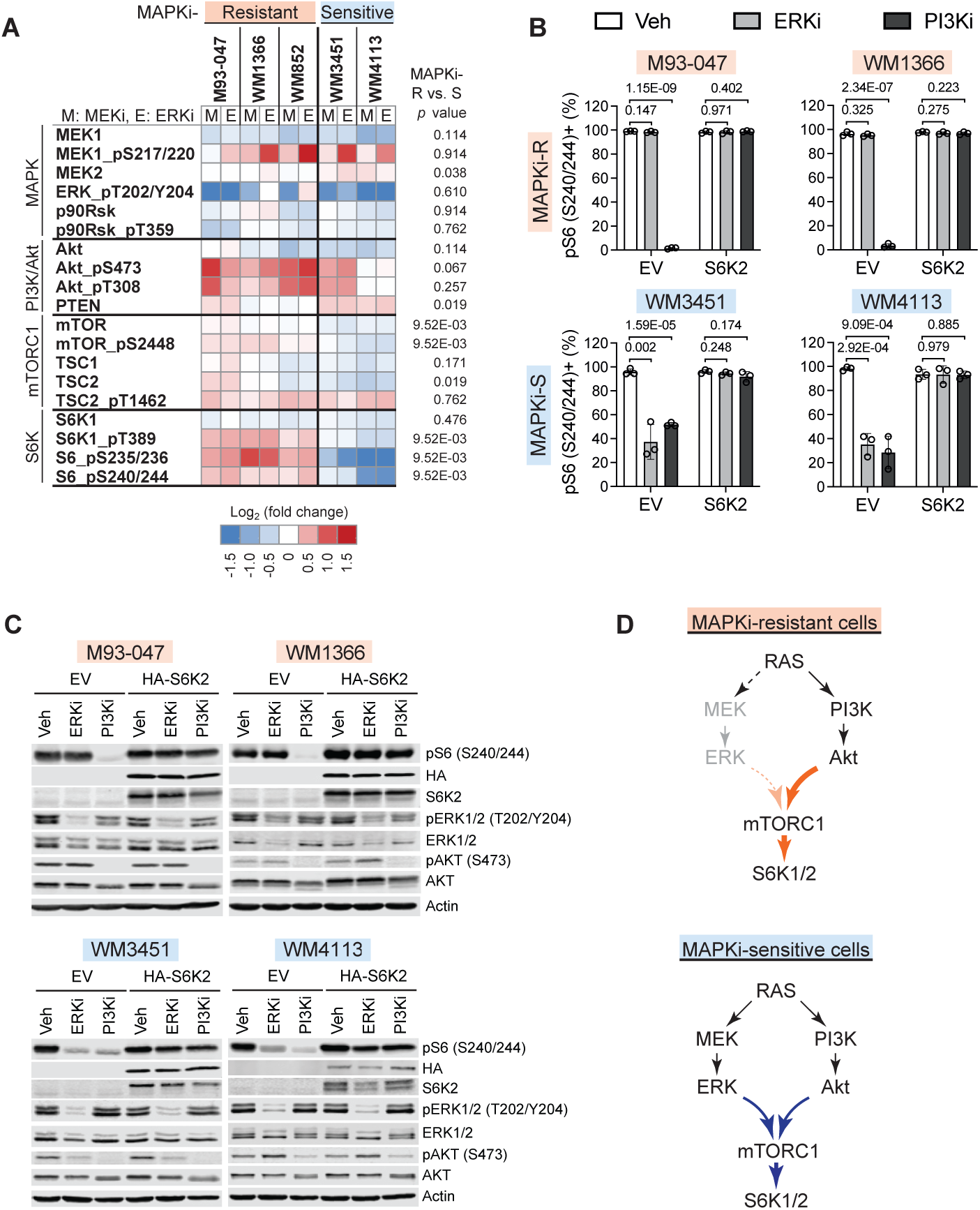
S6K is persistently activated in MAPKi-resistant NRAS mut melanoma. **A.** NRAS-mutant melanoma cells were treated for 48 h with the MEKi trametinib (M; 10 nM) or the ERKi SCH772984 (E; 100 nM) and changes in protein abundance or phosphorylation were determined by reverse phase protein array (RPPA) analysis (n = 3). Heatmap depicts log_2_ fold changes in MAPK, PI3K, mTOR, and S6K pathways relative to vehicle controls. Differences in response between resistant versus sensitive cell lines were calculated with unpaired, two-sided Mann-Whitney tests, indicating differential regulation of S6K signaling between MAPKi-resistant and -sensitive cell lines. **B**. MAPKi-resistant (M93-047, WM1366) or -sensitive (WM3451, WM4113) melanoma cells were transduced with doxycycline-inducible constitutively active HA-tagged S6K2T388E or empty vector (EV), pre-treated with 0.25 µg/ml doxycycline for 48 h, and then treated with DMSO vehicle (Veh), ERKi SCH984 (1 µM) or PI3K/mTORi GSK2126458 (GSK458, 100 nM) for an additional 24h. Percentage of pS6(S240/244)+ cells was determined by flow cytometry; bar graphs show mean ± SD (n = 3); *p*-values were calculated by unpaired, two-tailed Student’s t-tests. **C.** Cells treated as in (**B**) were probed by immunoblotting with the indicated antibodies. **D**. Diagram illustrating that in MAPKi-resistant NRAS-mutant melanoma cells, S6K is primarily regulated by the PI3K/mTOR pathway. In contrast, in MAPKi-sensitive cells, S6K relies on both the MAPK and the PI3K pathways.

Since S6K is also a bona fide effector of the PI3K/mTOR pathway(*27*), we further examined the impact of MAPK or PI3K/mTOR signaling on S6K. We ectopically expressed constitutively active S6K(*15*) (S6K1^T389E^ or S6K2^T388E^) constructs in MAPKi-sensitive or resistant cells and assessed levels of pS6^S240/244^, a site exclusively phosphorylated by S6K(*15*) (Fig. 1B-C and Fig.S1H-I). Treatment of MAPKi-S cells with MAPKi (SCH984) or PI3K/mTORi (GSK458) decreased phosphorylation of S6; ectopic expression of S6K1^T389E^ or S6K2^T388E^ rescued pS6 levels in cells treated with SCH984 or GSK458 (Fig. 1B-C and Fig. S1I). These data indicate that S6K mediates both MAPK and PI3K/mTOR signaling in MAPKi-S cells. In contrast, treatment of MAPKi-R cells with MAPKi had no effect on phosphorylation of S6, whereas treatment with PI3K/mTORi effectively suppressed pS6 (Fig. 1B-C and Fig. S1I). Ectopic expression of S6K1^T389E^ or S6K2^T388E^ rescued pS6 levels in MAPKi-R cells treated with PI3K/mTORi (Fig. 1B and Fig. S1I). Together, these data indicate that S6K activity is regulated by the PI3K/mTOR pathway in MAPKi-R cells. Moreover, these findings highlight that different upstream pathways differentially impinge on S6 kinase in MAPKi-R *vs.* MAPKi-S cells (Fig. 1D).

We next examined the expression of S6K1/2 in patient samples by interrogating the TCGA skin cutaneous melanoma dataset. We noted that S6K2 mRNA levels were inversely correlated with S6K1 (n=443, R=-0.47, p val=9.23 e-26; Fig. 2A). We also noted that in melanoma patients, high expression of S6K2 (but not S6K1) was associated with poor survival (Fig. 2B). Furthermore, analysis of RNA-seq data from NRAS^mut^ melanoma patient-derived cells(*28*) (Fig. 2C) revealed that S6K2 mRNA levels are significantly higher in MEKi-R tumor cells than in MEKi-S tumor cells. S6K1 mRNA levels were not significantly different in MEKi-S *vs.* MEKi-R-patient derived tumor cells. These data support the notion that S6K2 could have an important role in melanoma and MAPKi resistance, and thereby constitutes a vulnerability in this tumor type.

**Fig. 2.**
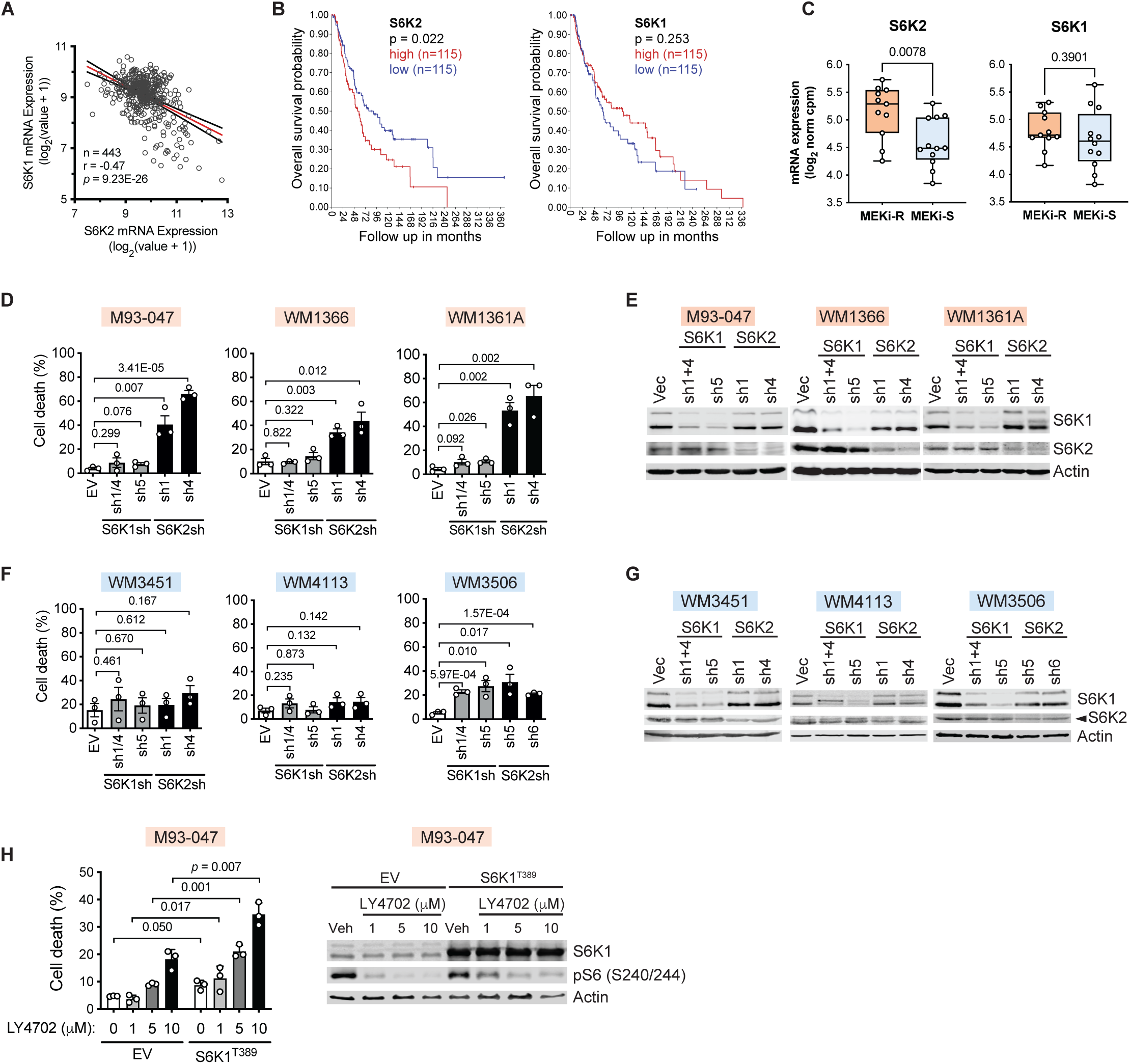
S6K2 is a vulnerability in NRAS-mutant melanoma resistant to MAPK inhibitors. **A.** S6K1 mRNA levels inversely correlate with S6K2 (443 patients; TCGA Skin Cutaneous Melanoma (SKCM); accessed via cBioPortal 5^th^ Feb 2022). **B**. Kaplan-Meier survival curve analysis was performed using R2: Genomics Analysis and Visualization. Overall survival curves are depicted for patients with high vs. low S6K2 or S6K1 expression (TCGA SKCM; n = 470; first *vs.* last quartile cut-off). **C.** Expression of S6K2 (*RPS6KB2*) and S6K1 (*RPS6KB1*) in trametinib-sensitive (MEKi-S; n=12) or -resistant (MEKi-R; n=11) patient-derived NRAS^mut^ melanoma cell lines as determined by RNA sequencing(*28*). Three MAPKi-R (**D**, **E**) or MAPKi-S (**F, G**) cell lines were transduced with lentiviruses encoding S6K1 or S6K2 shRNA and analyzed for cell death 7 days post infection (dpi) by Annexin V/PI (**D, F**) (mean ± SD; n=3; unpaired two-tailed Student’s *t*-test) or by immunoblotting (**E, G**) (n = 2). **H.** M93-047 cells were transduced with constitutively active S6K1^T389E^, treated with DMSO or the pan-S6K inhibitor LY2584702 (LY4702; 1, 5, 10 µM) for 48 hours and analyzed for cell death (left panel) or by immunoblotting (right panel).

To assess the role of S6K in NRAS-mutant melanoma cells, we silenced S6K1 or S6K2 in 6 NRAS-mutant melanoma lines (3 MAPKi-S and 3 MAPKi-R) using multiple hairpins. Depletion of S6K2, but not S6K1, induced cell death in MAPKi-R cells (Fig. 2D, E). Notably, neither depletion of S6K1 nor S6K2 triggered substantial cell death in MAPKi-S NRAS-mutant melanoma cells (Fig. 2F, G). These data suggest S6K2 is a vulnerability in NRAS-mutant melanoma cells that are resistant to MAPKi. To mimic the effects of an S6K2-specific inhibitor we ectopically expressed activated S6K1 in cells treated with the pan-S6K inhibitor LY2584702 (LY4702) (Fig. 2H). Activated S6K1 enhanced LY4702 cytotoxicity, rather than compensating for it, suggesting that sensitivity to pan-S6K inhibition may be mediated via S6K2 inhibition. Collectively, these results indicate that selective inhibition of S6K2, in the context of active S6K1, induces death of MAPKi-R NRAS-mutant melanoma cells.

### Depletion of S6K2 triggers lipid peroxidation and facilitates cell death

To investigate the mechanism(s) whereby S6K2 depletion triggers cell death in MAPKi-R melanoma cells, we surveyed the proteome to identify proteins that were differentially affected by depletion of S6K2 *vs.* S6K1 in MAPKi-R melanoma cells. Depletion of S6K2 was coupled to enhanced expression of proteins involved in lipid synthesis, fatty acid synthesis, uptake and activation, and phospholipid remodeling (Fig. 3A, Fig. S2A, and Supplementary Table 3). To further assess if lipid metabolism was perturbed by S6K2 depletion, we performed unbiased global lipidomic analysis in S6K2 or S6K1-depleted cells. Notably, depletion of S6K2 led to an enrichment of specific lipid species containing polyunsaturated fatty acyl chains (PUFAs) such as phosphatidylcholine, phosphatidylethanolamine and phosphatidylglycerol (Fig. 3B, Supplementary Table 4), consistent with disrupted lipid homeostasis. These results suggest that S6K2 blockade is coupled to an enhanced degree of fatty acid unsaturation in some lipid classes.

**Fig. 3.**
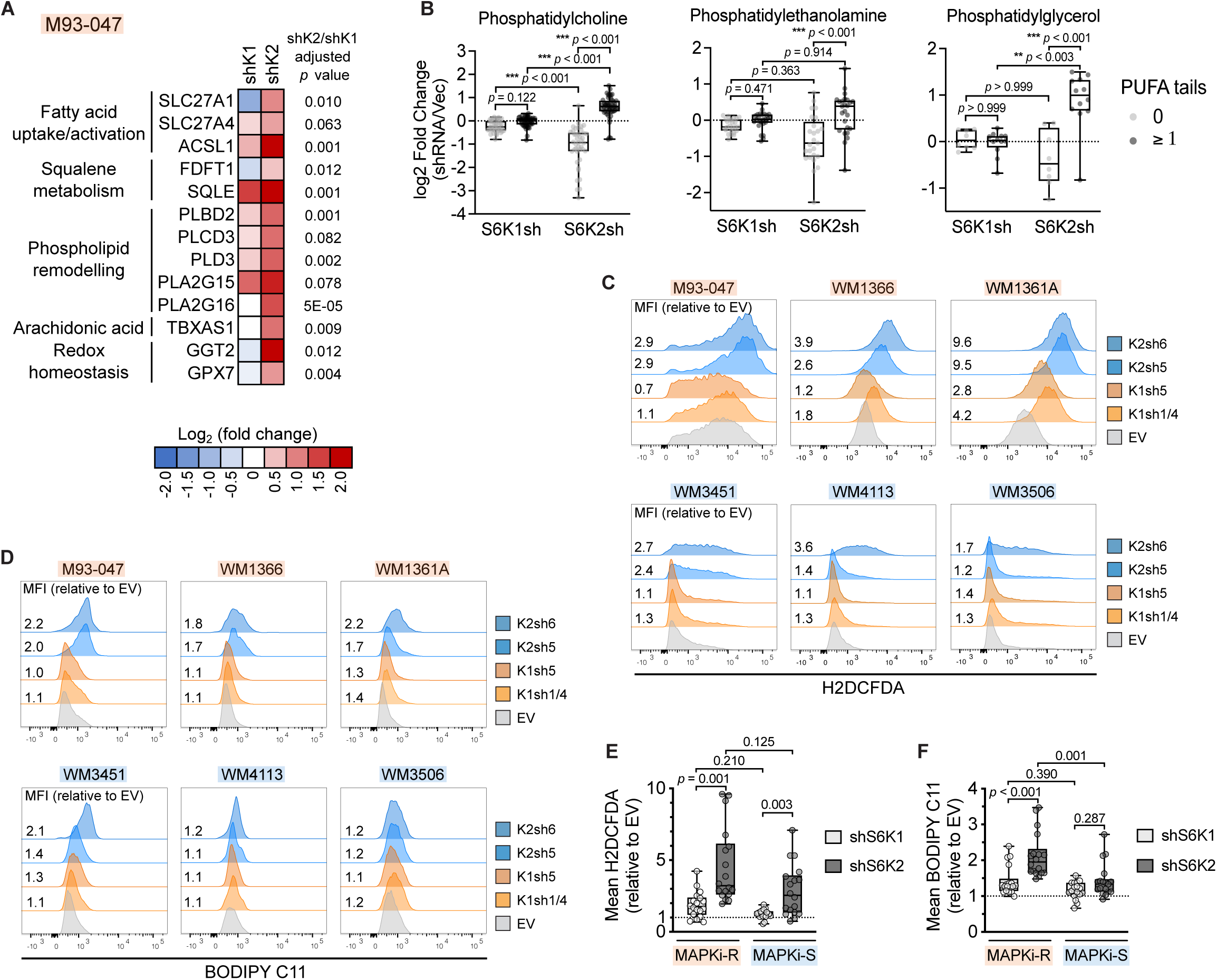
Depletion of S6K2 triggers lipid peroxidation. **A., B.** NRAS-mutant melanoma cells (M93-047) were transduced with lentivirus encoding S6K1 or S6K2 shRNA, or non-targeting vector control (Vec) and subjected to proteomic (**A**) or lipidomic (**B**) analysis (4 dpi). **A.** Heatmap depicting abundance of selected proteins involved in lipid metabolic pathways relative to vector control; *p*-values denote two-tailed Student’s *t*-tests of shS6K2 *vs.* shS6K1 adjusted for multiple hypothesis testing at a 10% FDR (Benjamini-Hochberg). **B.** Box and whiskers plot depicting classes of lipids differentially affected by shS6K2 *vs.* shS6K1. Median, upper quartile and lower quartiles are shown. Each dot indicates an individual lipid species. Details of lipids are presented in Supplementary Table 4. **C-F** Three MAPKi-R and three MAPKi-S cell lines were transduced with lentiviruses encoding S6K2 shRNA, S6K1 shRNA or empty vector control (EV). ROS levels (H2DCFDA; **C, E**) or lipid peroxidation (Bodipy-C11; **D**, **F**) were assessed 5 days post infection by flow cytometry. **C, D.** Representative histograms of three independent experiments (n=3) showing mean fluorescence intensity relative to EV control. **E, F** Box and whisker plots of pooled MAPKi-R/S data including 3 MAPKi-R and 3 MAPKi-S cell lines transduced with two different shRNAs for S6K1 or S6K2. Median, upper quartile and lower quartile are shown. **B, E, F** Data are from three independent biological replicates. Kruskal-Wallis ANOVA tests were used to compare groups and corrected for multiple comparisons with Dunn’s test.

We also noted that depletion of S6K1 or S6K2 led to increased levels of proteins involved in the oxidative stress response, including ROS sensing/detoxifying enzymes such as GGT2, SOD2 and PARK7 (Fig. 3A, Fig. S2A-B and Supplementary Table 3). While depletion of either S6K1 or S6K2 (Fig. S2H) led to increased ROS levels (Fig. 3C, E), depletion of S6K2 induced significantly higher relative ROS levels compared to S6K1 depletion in both MAPKi-R cells and MAPKi-S cells (Fig. 3C, E). Although depletion of S6K2 increased ROS levels in MAPKi-S cells, these cells did not undergo cell death (Fig. 2F), implying that ROS alone are not sufficient to induce cell death or that MAPKi-S cells are able to counteract redox imbalance. The perturbation of redox and lipid homeostasis upon S6K2 knockdown raised the possibility that S6K2 depletion could be triggering lipid peroxidation(*29*). Indeed, depletion of S6K2 (but not S6K1) substantially enhanced lipid peroxidation and oxidative stress, as indicated by the lipid ROS sensor Bodipy-C11 (Fig. 3D, F), generation of 4-HNE adducts (Fig. S2C, E), and increased levels of nucleic acid oxidative damage (8-OHdG; Fig. S2D, F) in MAPKi-R cells. Whereas general S6K2 depletion induced similar increase in general ROS levels in both MAPKi-S and MAPKi-R cells, lipid ROS were preferentially induced in MAPKi-R cells (Fig. 3C-F). Together, these results support the premise that depletion of S6K2 enhances lipid metabolism and ROS levels, which jointly trigger lethal lipid peroxidation and oxidative cell death in MAPKi-R NRAS-mutant melanoma.

### S6K2 depletion activates a terminal unfolded protein response (UPR) with concomitant upregulation of PPARα

Excessive lipid peroxidation has been linked to different types of cell death including apoptosis and ferroptosis(*29–35*). To identify S6K2 effectors that could trigger cytotoxicity, we assessed global transcriptional changes induced by S6K2 depletion. Transcriptomic analysis indicated a general enrichment of unfolded protein response (UPR) regulators in both S6K1- and S6K2-depleted cells, whereas XBP-1 and target genes of the lipid metabolism regulator PPAR(*36, 37*) were selectively increased in S6K2-depleted cells (Fig. 4A, G, and Supplementary Table 5). Oxidative or metabolic stress can disrupt endoplasmic reticulum (ER) homeostasis, promoting the aberrant accumulation of misfolded proteins, inducing ER stress and further dysfunction of lipid metabolism(*38*). ER stress initiates the unfolded protein response (UPR) through BiP-mediated activation of three canonical pathways: IRE1α-XBP-1, PERK-CHOP and ATF6(*39*), either restoring ER homeostasis under conditions of mild ER stress or triggering apoptosis when ER stress is severe and persistent. We found increased levels of activated, spliced XBP-1 and its regulator phospho-IRE1α in S6K2-depleted cells (Fig. 4A-C). In contrast, S6K2 depletion had a negligible effect on PERK-CHOP signaling (Fig. 4B). Treatment of MAPKi-R NRAS-mutant melanoma cells with the IRE1α inhibitor Kira6 partially attenuated cell death induced by S6K2 depletion, indicating that the cytotoxic effects of S6K2 depletion are partially mediated by a terminal UPR that activates apoptosis. Additionally, treatment with the pan-caspase inhibitor zVAD or the antioxidant glutathione (GSH) also attenuated cell death (Fig. 4D). In contrast, treatment of MAPKi-R NRAS-mutant melanoma cells with the ferroptosis inhibitor, ferrostatin-1, did not attenuate cell death triggered by S6K2 depletion (Fig. S3A). Upregulation of XBP-1 splicing and PPARα expression caused by S6K2 knockdown was suppressed by IRE1α inhibition with Kira6 in a dose-dependent manner, suggesting that PPARα could be a key effector downstream of ER stress (Fig. 4E, Fig. S3B). Since XBP-1 directly binds to and activates PPARα and PPARγ(*40, 41*), we assessed the expression of the PPAR gene family following S6K2 knockdown (Fig. 4F, Fig. S3C). PPARα expression was consistently elevated by S6K2 depletion in MAPKi-R cells, whereas PPARγ was upregulated in a single cell line. Further, S6K2 depletion led to upregulation of representative PPAR targets involved in lipid metabolism in both MAPKi-R cell lines (Fig. 4G-H). Together, these data suggest that S6K2 depletion causes ER stress and that the IRE1α-XBP-1-PPARα signaling network contributes to cell death and lipid imbalance triggered by S6K2 depletion.

**Fig. 4.**
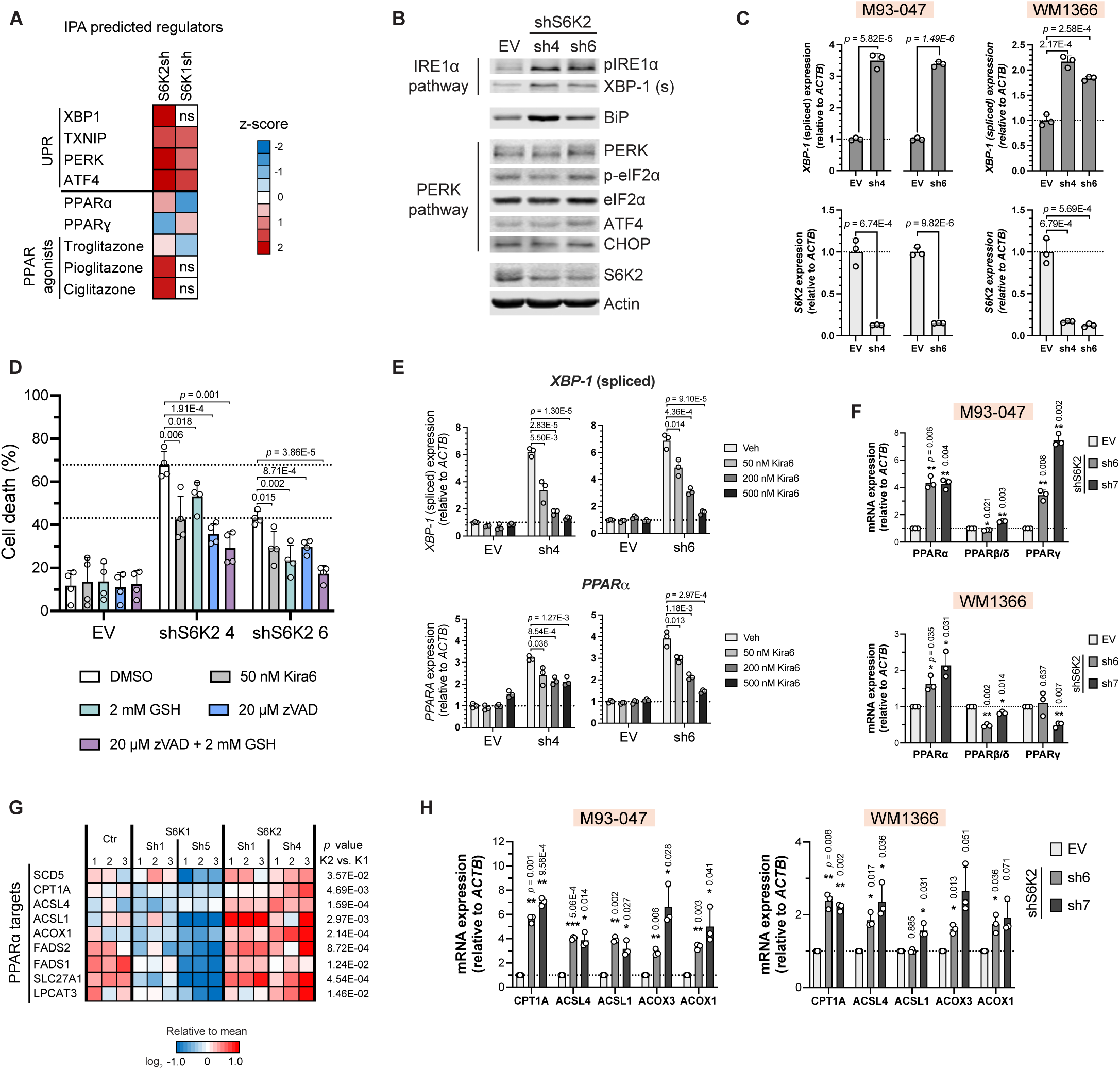
S6K2 depletion activates a terminal unfolded protein response (UPR) and PPARα. MAPKi-R were transduced with S6K1 shRNA, S6K2 shRNA or vector control. **A.** Cells were analyzed by RNA sequencing (3 dpi). Genes significantly altered by S6K1/2 depletion were analyzed by IPA. Heatmap depicts *z*-scores for predicted activity of unfolded protein response (UPR) regulators, PPAR genes and PPAR modulators. Positive/red: activated; negative/blue: inhibited by S6K1 or S6K2 shRNA; ns: no significant enrichment. **B., C.** Endoplasmic reticulum (ER) stress markers were monitored following transduction with shS6K2 or EV control by Western blotting (**B;** M93-047, 3 dpi) and qRT-PCR (**C**; M93-047, sh4 3 dpi, sh6 4 dpi; WM1366 4 dpi). **D.** M93-047 cells transduced with S6K2 shRNA or empty vector (EV) control were treated with the indicated compounds. Cell death was analyzed by Annexin V/PI staining 5 dpi (mean ± SD (n=4); unpaired two-sided *t*-test). **E.** M93-047 cells transduced with EV/shS6K2 were treated with the IRE1α inhibitor Kira6. Levels of spliced XBP-1 and PPARα were quantified by qRT-PCR (3 dpi, sh4 or 4 dpi, sh6). **F, H** PPAR gene family (**F**) and PPAR targets (**H**) mRNA expression were determined by qRT-PCR 4 dpi with S6K2 shRNA/EV. **G.** M93-047 cells were transduced with EV, S6K1 or S6K2 shRNA (3 dpi) and analyzed by RNA sequencing. Heatmap depicts mRNA expression levels of selected PPAR targets relative to the mean across samples; *p* value: Student’s *t*-test. **C, E, F, H.** Data are shown as mean ± SD from three independent experiments. \**p* ≤ 0.05, \*\**p* ≤ 0.01, \*\*\**p* ≤ 0.001; unpaired two-sided *t*-tests.

### S6K2 negatively regulates PPARα

We next sought to define the mechanism whereby S6K2 could be regulating PPARα. The transcriptional activity of PPARα is inhibited by the nuclear receptor corepressor NCoR1(*40*), and S6K2 has been implicated in this regulation(*41*). Hence, we examined whether S6K2 could regulate this interaction in NRAS-mutant melanoma cells using the technique of proximity ligation assays (PLA). PLA supported the existence of a complex between S6K2, PPARα and its co-repressor NCoR1 in MAPKi-R cells (Fig. 5A-C). S6K2 depletion markedly diminished the interaction of NCoR1 with PPARα in MAPKi-R cells (Fig. 5D, F, Fig. S4A), indicating that S6K2 blockade is coupled to PPARα activation. In comparison, we noted a weak and diffuse PPARα-NCOR1 interaction in MAPKi-S cells, which was slightly enhanced following S6K2 depletion (Fig 5E, F, Fig. S4B). Together, these data support a model whereby S6K2 regulates the interaction between of NCoR1 and PPARα; S6K2 depletion disrupts this complex leading to activation of the PPARα axis, thereby facilitating transcription of genes encoding proteins involved in lipid peroxidation and oxidative cell death in MAPKi-R NRAS-mutant melanoma.

**Fig. 5.**
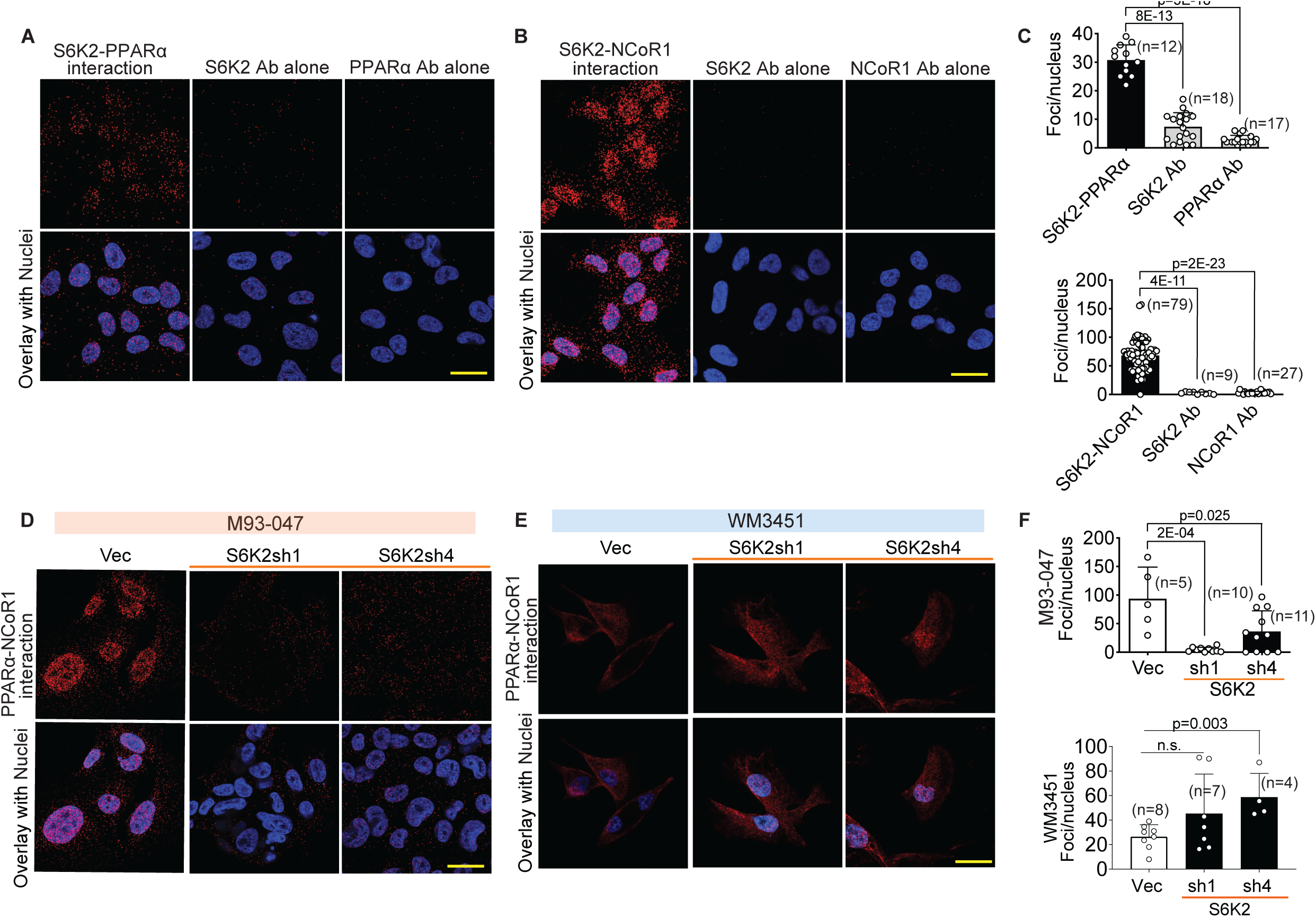
S6K2 negatively regulates PPARα. **A., B.** Interaction of S6K2 with PPARα or NCoR1 was determined by proximity ligation assay (PLA). Representative images. Scale bar = 30 µM. **C.** Bar graphs show quantification of PLA signals for a (top) and b (bottom); mean ± SD (n denotes fields quantified). S6K2 Ab, PPARα Ab or NCoR1 Ab indicates single antibody controls. **D, E.** Transduced cells (4dpi) were analyzed by PLA. (**D**) MAPKi-R M93047 cells, (**E**) MAPKi-S WM3451 cells. Representative images shown, Scale bar = 30 µM. **F.** Bar graphs show quantification of PLA signals for d (top) and e (bottom); mean ± SD (n denotes fields quantified).

### PUFAs potentiate the oxidative cell death inducing effect of PPARα agonists

Based on our data indicating that PPARα facilitates lethal lipid peroxidation, we next asked if PPARα agonists could induce anti-melanoma effects. We noted that PPARα protein levels were higher in MAPKi-R NRAS-mutant melanoma cells (Fig. 6A and Fig. S5A) and that MAPKi-R cells were more sensitive to the PPARα agonist fenofibrate (FNB) than MAPKi-S/PPARα-low cells (Fig. 6B and Fig. S5B). FNB is a weak PPARα agonist, which can decrease GSH levels(*42*) and has been shown to modify lipid and lipoprotein composition and metabolism by a variety of mechanisms(*43*). Treatment with FNB alone did not induce cell death in MAPKi-R cells (Fig. 6C). Since our lipidomic data revealed S6K2 depletion induces accumulation of PUFAs (Fig. 3B), we wondered whether addition of docosahexaenoic acid (DHA) as a source of PUFAs could potentiate the effect of FNB in inducing lethal lipid peroxidation and cell death. To this end, we treated MAPKi-R and MAPKi-S cells with FNB +/-DHA. While neither FNB nor DHA as single agents induced significant cell death, the combination of FNB + DHA triggered significant cell death in MAPKi-R cells, but not in MAPKi-S NRAS-mutant melanoma cells (Fig. 6C). Similar to S6K2 depletion, the FNB/DHA combination induced ROS levels (Fig. S5C). Consistent with the effects of S6K2 depletion, treatment with FNB+DHA induced lipid peroxidation selectively in MAPKi-R cells (Fig. 6D, E). Further supporting the notion that the FNB/DHA combination phenocopies S6K2 depletion, the antioxidant GSH, pan-caspase inhibitor, zVAD and IRE1α inhibitor, Kira6 attenuated FNB/DHA-induced cell death (Fig 6F, G). Additionally, the FNB/DHA combination induced both spliced XBP-1 and PPARα (Fig. S5D, Fig 6H, I), and cotreatment with Kira6 suppressed the activation of XBP-1. Moreover, the FNB/DHA combination induced cell death in MAPKi-R 3D melanoma spheroids (Fig 6J). Taken together, these results indicate that MAPKi-R melanoma cells are sensitive to pharmacological activation of PPARα in combination with enhanced fatty acid desaturation and lipid peroxidation.

**Fig. 6.**
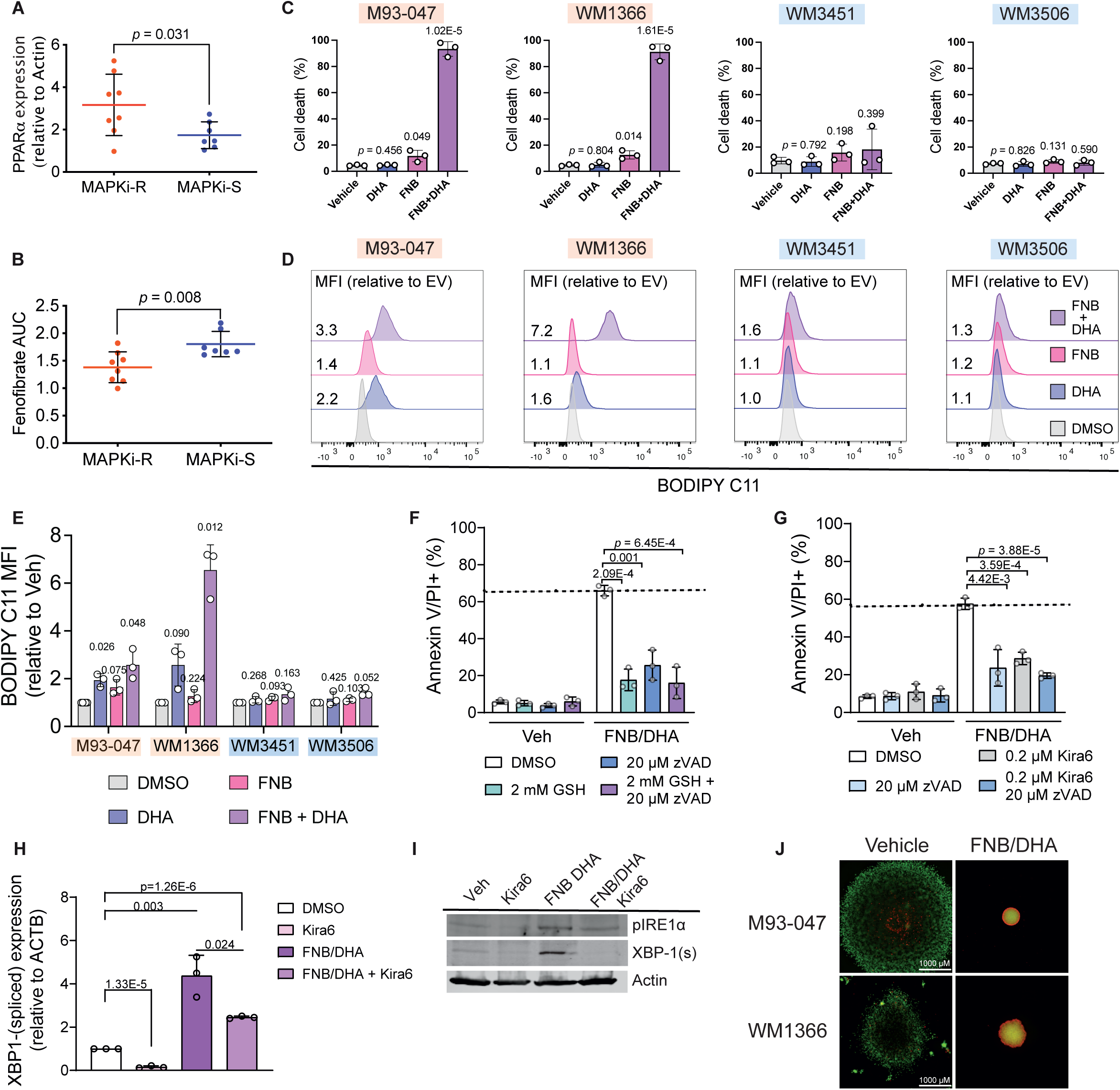
PUFAs potentiate the oxidative cell death inducing effect of PPARα agonists. **A.** Relative expression of PPARα in MAPKi-R (red, n=?) and MAPKi-S (blue, n=?) cells determined by immunoblotting (see Fig. S6A), was compared by Student’s T-test. Data represent average PPARα levels from two independent experiments. **B.** AUC for MAPKi-R and MAPKi-S cells (shown in Fig. S6c) were compared by Student’s T-test. **C.** Cells were treated with 50 µM fenofibrate, 7.5 µM DHA, as single agents or in combination for 72 h. Cell death were assessed by Annexin V/PI positivity (mean ± SD; n=3; unpaired two-tailed Students *t*-test). **D., E.** Cells were treated with 50 µM fenofibrate, 7.5 µM DHA, as single agents or in combination. Lipid peroxidation was assessed by Bodipy-C11. (**D**) representative histograms and (**E**) data from three independent experiments showing mean fluorescence intensity relative to vehicle control. **F., G.** M93-047 cells were pre-treated with the indicated compounds [(**F**) antioxidant GSH (2mM), pan-caspase inhibitor (20 µM zVAD) for 24h or (**G**) pan-caspase inhibitior (20 µM zVAD), UPR/IRE1 inhibitor 0.2 µM Kira6 for 48h)] and then treated with 50 µM fenofibrate, and/or 7.5 µM DHA for 24h. Cell death was analyzed by AnnexinV/PI positivity. Graphs show mean ± SD (n=3), individual dots represent independent experiments; unpaired two-sided *t*-test). **H., I.** M93-047 cells were treated with 0.2 µM Kira6 for 48h and then treated with 50 µM fenofibrate, 7.5 µM DHA for 24h. (**H-I**) Levels of spliced XBP-1 were quantified by qRT-PCR, mean ± SD (n=3); unpaired two -sided *t*-test or Western blotting (**I**). **J.** MAPKi-R M93-047 and WM1366 cells grown as collagen-embedded 3D spheroids were treated with Vehicle (left) 50 µM fenofibrate, 7.5 µM DHA (right) for 5 days. Spheroids were stained with Calcein (AM) (green; live cells) and EtBr (red; dead cells) and imaged with a fluorescence microscope. Representative images of three replicates are shown; the scale bar represents 1000µM.

### MAPKi-resistant NRAS-mutant melanoma is sensitive to the combination of PPARα agonists and PUFAs

We next evaluated the efficacy of combining PPARα agonists and DHA *in vivo*. Combining FNB with DHA suppressed the growth of established NRAS-mutant MAPKi-R xenografts (Fig. 7A) and syngeneic tumors (Fig. 7B)(*44, 45*) with no appreciable toxicity (Fig. S6A). Furthermore, FNB+DHA significantly increased the survival of mice bearing MAPKi-R PDXs (Fig. 7C, D, Fig. S6B). Importantly, FNB+DHA induced substantial lipid peroxidation and oxidative damage in MAPKi-R xenografts and PDXs (Fig. 8A-D). These results provide proof-of-principle that combining PPARα agonists with PUFAs facilitates lipid peroxidation, elicits anti-tumor activity, and restrains MAPKi-R NRAS-mutant melanoma. By selectively blocking S6K2 or harnessing the S6K2 subnetwork, we have identified a strategy that could be potentially exploited as an anti-tumor approach in NRAS-mutant melanoma resistant to MAPKi.

**Fig. 7.**
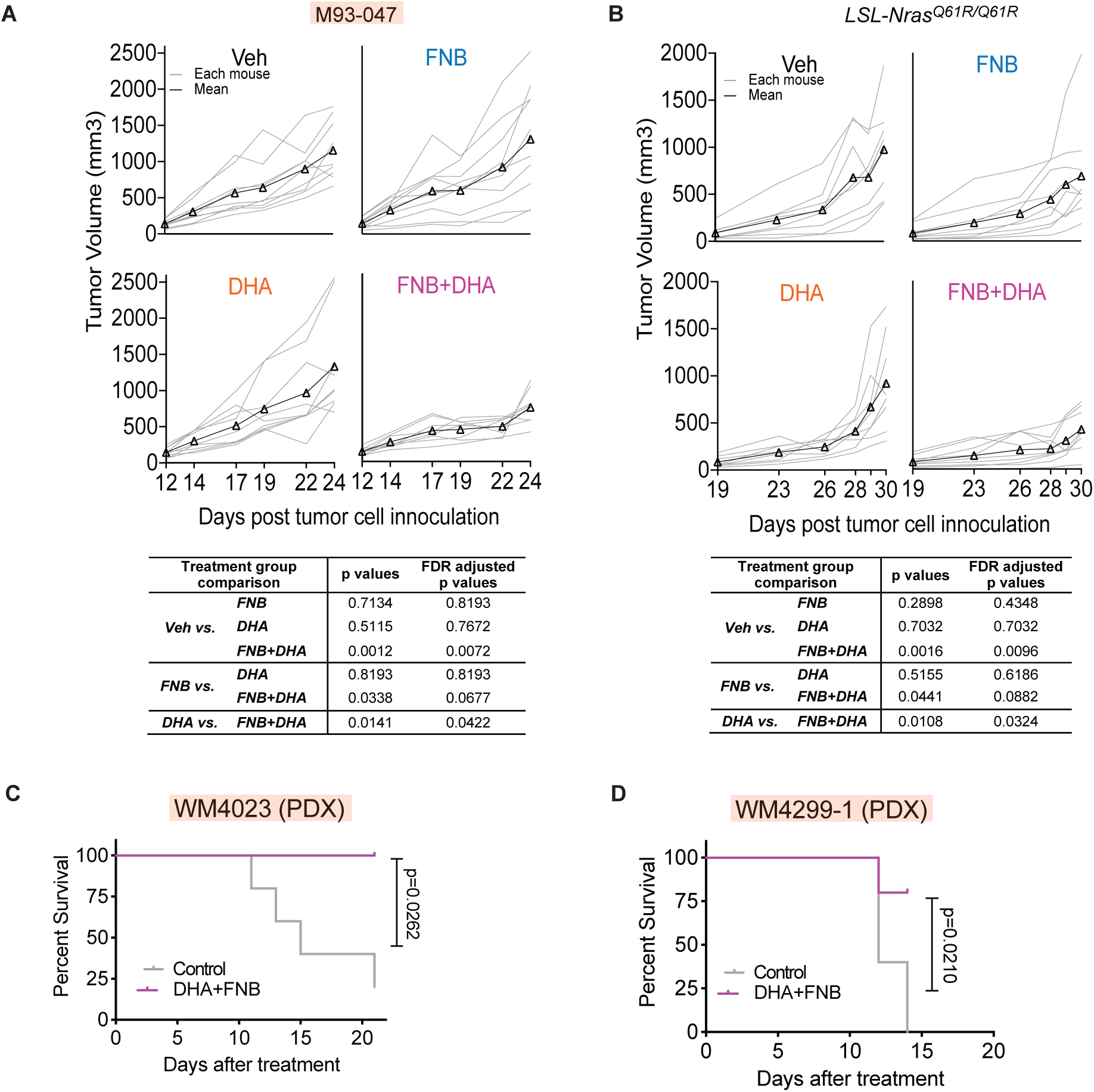
MAPKi-resistant NRAS-mutant melanoma is sensitive to the combination of PPARα agonists and PUFA. **A.** Mice bearing MAPKi-R M93-47-derived subcutaneous tumors were treated with fenofibrate (200 mg/kg), DHA (300 mg/kg) as single agents or in combination. Tumor volume for individual mice (gray lines) or mean (black lines, n=8) is shown. **B.** C57BI/6j mice bearing MAPKi-R *LSL-Nras^Q61R/Q61R^ (TpN^61R/61R^)* syngeneic tumors were treated with vehicle (n=8), fenofibrate (200 mg/kg, n=9), DHA (300 mg/kg, n=8) or fenofibrate + DHA combination (n=8) when tumors reached ∼85 mm^3^ (19 dpi). **A-B** p-values and FDR adjusted p-values were estimated from a linear mixed-effect model with all follow-up subjects and time points. **C., D.** Mice bearing PDXs were fed control chow or DHA (0.15% w/w) plus FNB (0.15% w/w) laced chow when tumors reached ∼100-150 mm^3^ (MAPKi-R WM4023, control n=5, DHA+FNB n=4) or (MAPKi-S WM4299-1, control n=5, DHA+FNB n=5). Mice were followed until tumors reached a pre-defined volume (1200 mm^3^). P values were calculated using log-rank (Mantel-Cox) test.

**Fig. 8.**
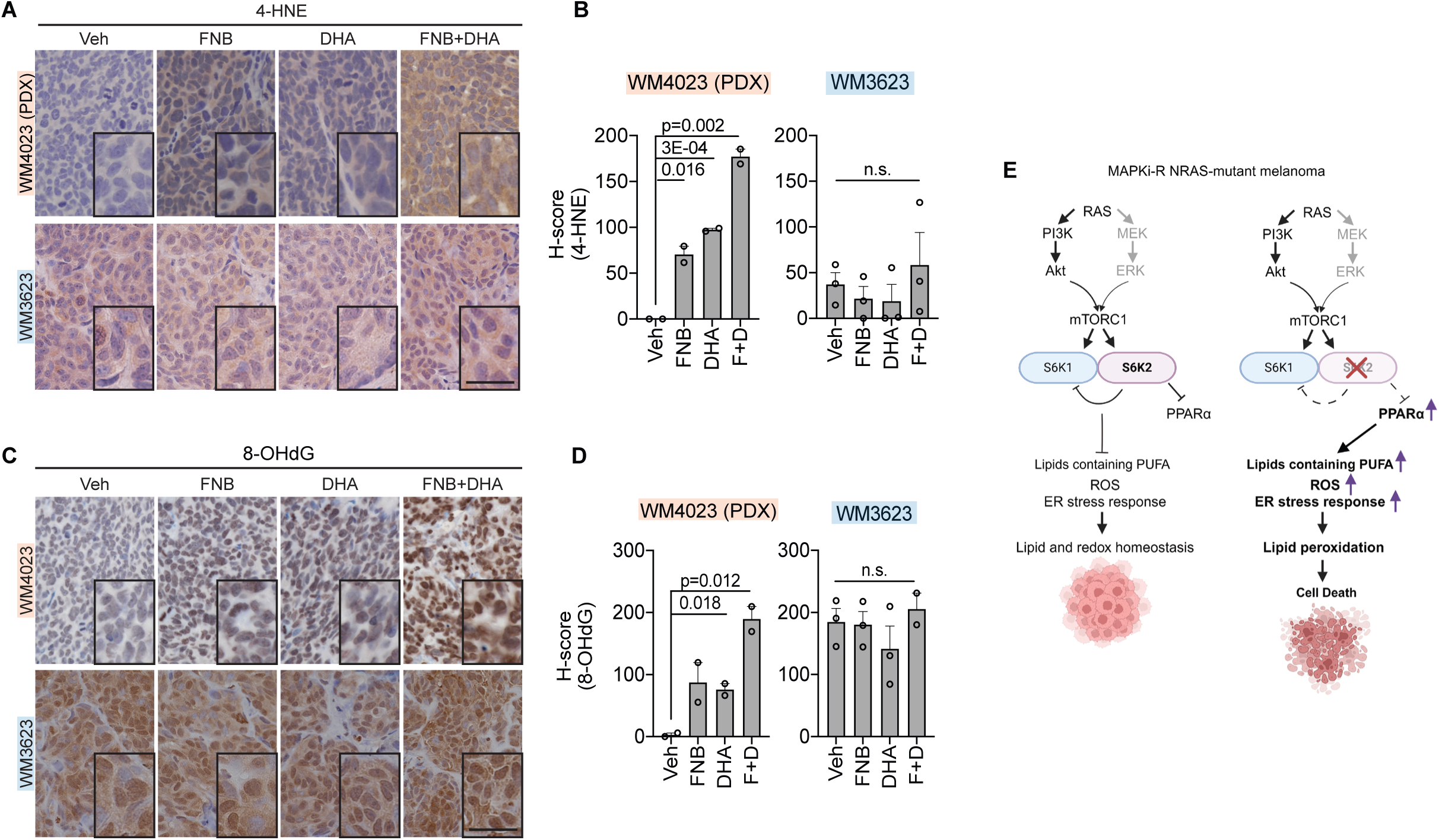
PPARα agonists and PUFA induce lipid peroxidation and oxidative damage. Tumors derived from Mice bearing MAPKi-R WM4023 (PDX), or MAPKi-S WM3623 xenografts treated with fenofibrate (200 mg/kg) or DHA (300 mg/kg) as single agents or in combination for 7 days were analyzed by immunohistochemistry for (**A**) 4-HNE (**C**) 8-OHdG. **A., C.** Representative images. Scale bar = 25 mm. **B., D.** Tumors were scored using QuPath software. Bar graphs show H-score from representative fields (Mean ± SD; WM4023 n=2, WM3623 n=3) *t*-test; n.s., not significant. **E.** Graphical model (created with BioRender.com) depicting MAPKi-R NRAS-mutant melanoma when S6K2 active and present (left panel) and when S6K2 is depleted (right panel).

## DISCUSSION

Despite significant advances in treating melanoma, effective therapies for NRAS-mutant tumors are sorely needed. One approach to inhibiting NRAS that has been investigated is targeting its proximal downstream effectors, mainly the MAPK and PI3K pathways. While inhibiting either pathway alone is barely effective in NRAS-mutant tumors, inhibiting both pathways leads to unacceptable toxicities in patients. We hypothesized that identifying and inhibiting convergent subnetworks, within the MAPK and PI3K super-networks, could spare the toxicities while triggering death of cancer cells. We therefore performed a deep analysis of the signaling networks in MAPKi-resistant NRAS-mutant melanoma. By comparing the effects of MAPK inhibition in MAPKi-S *vs.* MAPKi-R NRAS-mutant melanoma, we mapped a critical subnetwork downstream from the convergence of PI3K and MAPK. We found that a S6 kinase 1/2 (S6K1/2)-PPARα subnetwork is vital for cancer lipid metabolism. We uncovered that S6K1/2 is regulated by both the MAPK and PI3K pathways in MAPKi-S cells; in contrast, S6K1/2 is regulated by PI3K in MAPKi-R NRAS-mutant melanoma. Further, we show that uncoupling S6K1 and S6K2, by selective S6K2 depletion, triggers a lipid metabolic imbalance featuring ER stress and lipid peroxidation, which leads to cell death preferentially in MAPKi-R NRAS-mutant melanomas (Fig. 8E). Both the IRE1α-XBP-1 and PERK-CHOP pathways are implicated in initiating ER stress-induced apoptosis(*46*). Upon S6K2 knockdown, ER stress predominantly activated the IRE1α-XBP-1 pathway. Moreover, cell death was suppressed by the IRE1α inhibitor Kira6, indicating that IRE1α-XBP-1 signaling plays a key role in mediating cell death elicited by S6K2 suppression. Our findings underscore that MAPKi-resistant and -sensitive cells have different signaling dependencies both upstream and downstream of mTORC1 and illustrate specialized functions of S6K isoforms, highlighting the biological importance of isoform differentiation. Moreover, the identified subnetwork provides a strategy to pharmacologically induce lethal lipid peroxidation *in vivo*, which has been limited by the lack of compounds with sufficient bioavailability(*33, 47–49*). Of note, S6K2 KO mice are viable and develop normally(*21*), indicating that S6K2 is not an essential gene. Collectively, these data suggest that selective targeting of S6K2 or its effectors represents a viable opportunity for therapeutic intervention in S6K2-dependent tumors such as MAPKi-R NRAS-mutant melanoma.

Our study establishes that selectively depleting S6K2 induces ER stress and lethal lipid peroxidation in MAPKi-R NRAS-mutant melanomas. This suggests that selective inhibition of S6K2 could curb melanoma through a mechanism unachievable with PI3K or mTOR inhibitors, which inhibit both S6K1 and S6K2. However, selective S6K2 inhibition has not been explored in pre-clinical melanoma models and only one S6K2-selective chemical probe has been reported(*50*), in part due to the high sequence similarity of S6K2 with S6K1 and the lack of S6K2 crystal structure(*18, 24*). We foresee that our findings will spur the development of potent, bioavailable S6K2 inhibitors. Meanwhile, we have identified actionable targets whose modulation could be a surrogate for S6K2 inhibition and provide proof-of-concept for this strategy. The combination of FNB (PPARα agonist) plus DHA (PUFA) recapitulated the activation of ER stress and XBP-1-PPARα signaling observed with genetic depletion of S6K2, promoting lipid peroxidation and cell death. Importantly, both FNB and DHA are compounds that are used in humans with manageable toxicity profiles and, therefore, could be incorporated into new treatments for melanoma. Of note, MAPKi-R and MAPKi-S NRAS-mutant melanomas exhibit different levels of PPARα, which correlate with sensitivity to MAPKi and PPARα agonists. This provides a context in which our proposed approach would be most effective.

Previous studies from our group and others have linked enhanced or persistent mTORC1/S6K1 signaling with acquired resistance to MAPKi in BRAF-mutant melanoma(*7, 15, 51*) and to MEKi/Cdk4i in NRAS-mutant melanoma(*52, 53*). Inhibition of the mTORC1/S6K1 axis restrains MAPK-dependent melanomas and tumors with acquired resistance to MAPKi and/or CDK4i. In contrast to strategies relying on mTOR1/S6K1 inhibition, we exploited active lipid metabolism to induce cell death(*54*) by disrupting the S6K1/S6K2/PPARα subnetwork through selective S6K2 blockade. We noted that while S6K2 blockade led to enhanced expression of proteins involved in lipid synthesis, and accumulation of PUFAS, inhibition of S6K1 minimally perturbed lipid homeostasis. These data support the notion that selective S6K2 inhibition is required to induce lethal peroxidation in melanoma. Inducing severe ER stress and lethal lipid peroxidation provides a therapeutic opportunity to offset drug resistance. Previous studies have shown that MAPKi-R melanomas display elevated levels of oxidative stress and are vulnerable to compounds that enhance ER stress(*55, 56*). Moreover, activation of the UPR can promote anti-tumor immunity in a context-dependent manner(*57, 58*). Likewise, tumors characterized by distinctive metabolic states such as those displaying enhanced PUFA biosynthesis are innately vulnerable to lipid peroxidation(*59, 60*). In sum, our studies establish S6K2 and its effectors as a vulnerability in NRAS-mutant melanomas that are resistant to MAPKi. Our results underscore the significance of oncogenic NRAS-induced metabolic dependency and the role of S6K2 in supporting this addiction. We propose that harnessing this S6K2-dependent addiction could be exploited as a potential strategy to combat MAPKi-resistant NRAS-mutant melanoma, a highly aggressive tumor type with limited treatment options.

## ACKNOWLEDGMENTS

We would like to thank Shared Core Facilities at the Wistar Institute (Genomics, Proteomics and Metabolomics, Molecular Screening and Protein Expression, Histotechnology, Imaging, Flow Cytometry and Animal Facilities) for technical support. Support for the above shared resources was provided by Cancer Center Support Grant (CCSG) P30CA010815 to the Wistar Institute. The Thermo Q-Exactive HF-X mass spectrometer was purchased with NIH grant S10 OD023586. Dr. Yardena Samuels (Weizmann Institute of Sciences) shared RNA-seq data(*28*), Dr. Michael Weber provided essential reagents, cell lines and PDX were generous gift of Dr. M. Herlyn (The Wistar Institute). The RPPA analysis was performed by the MDACC RPPA core facility with support for shared resources provided by Cancer Center Support Grant (CCSG) CA016672 to MDACC. The Thermo Q-Exactive HF-X mass spectrometer was purchased with NIH grant S10 OD023586. Support for RPPA was provided by the Dr. Miriam and Sheldon G. Adelson Medical Research Foundation.

Work in our laboratory has been partially supported by NIH grants P01CA114046 (JV, MM), P50CA174523 (JV, ARG), R01CA268510 (JV), U54CA224070 (JV), DoD HT94252310914, The Melanoma Research Alliance and The V Foundation for Cancer Research (JV), the PA Department of Health (JV), The Wistar Science Accelerator Award and Goldblum Family Healthcare Fund (JV). DZ was supported by NIH pre-doctoral training grant T32 GM008275, BL were supported by NCI NRSA T32 CA009171 Cancer Biology Training Grant to the Wistar Institute. We dedicate this work to the memory of our friend and colleague Michael J. Weber.

## AUTHOR CONTRIBUTIONS

BL, ANG, H-YC, JV, MEM and DWS conceived and designed the study. BL, ANG, H-YC, and JV wrote the manuscript. BL, ANG, H-YC, SDA, ARG, DZ, PIR-U, SB, performed experiments, collected and analyzed data. XY and QL performed bio-statistical analysis. AVK performed bioinformatics analysis. YL and GM performed RPPA. All authors contributed, reviewed, and approved the manuscript.

## METHODS

### Cell culture

Melanoma cell lines and 293T cells were grown in RPMI1640 supplemented with 5% and 10% FBS respectively. FBS was purchased from Tissue Culture Biologicals (#101; Lot#170708, 105094, 101258, 101929) or Clontech (#631107; Lot#A301117007).

### 3D melanoma spheroids

Melanoma spheroids were generated and stained for live/dead cells as previously described (*45*) with the following modifications: spheroids were embedded into a collagen mixture (10% RPMI, 1.5 mg/ml collagen, 10% FBS, and 7.5% NaHCO_3_) and treated with DMSO or 50 µM Fenofibrate + 7.5 µM DHA.

### Source of compounds and chemicals

Cayman chemical: BVD-523 (#18298), Docosahexaenoic Acid (DHA; Cayman Chemical #90310 & MedChemExpress #HY-B2167/CS-6261), Fenofibrate (#10005368), GSH (#10007461), GSK2126458 (#17377), Kira6 (#19151), MEK-162 (#16996), SCH772984 (#19166), zVAD (zVAD(OMe)-FMK, #14463).

Selleckchem: Ferrostatin-1 (#S7243), LY2584702 (#S7704), Trametinib (#S2673).

Sigma-Aldrich: BrdU (5-Bromo-2’-deoxyuridine, #B5002), G418 (Sigma-Aldrich #G418-RO), PEG 300 (#202371), Propidium iodide (#P4864), RNase A (#R4642), Soybean oil (#S7381), TES (#T1375), Tween-80 (#P1754).

BODIPY™ 581/591 C11 (Thermo Fisher Scientific, #D3861), Doxycycline HCl (Research Products International Crop. #D43020), MTT (BIOSYNTH #T-3450), PSVue 643 (Molecular Targeting Technologies P-1006), Puromycin (Gemini Bio Products, #400-128P).

### Animal Studies

All animal studies were approved by the Institutional Animal Care and Use Committee (IACUC) at the Wistar Institute and performed in accordance with institutional guidelines. Mice were housed in groups of five/cage in an Association for the Assessment and Accreditation of Laboratory Animal Care (AAALAC) certified facility. NOD/LtScidIL2Rγ-null (NSG) mice (4 to 8-week-old, ∼ 25-30 g) were procured from The Wistar Institute. C57Bl/6 mice (4-week-old, ∼25 g) were obtained from Charles River.

For *in vivo* xenograft studies, 1 x 10^6^ cells were resuspended in 1:1 RPMI1640/Matrigel (Matrigel Matrix, Corning #354230) and implanted subcutaneously into the flanks of male NSG mice. Patient-derived xenografts were minced, mixed with Matrigel and implanted into male NSG mice.

For the syngeneic mouse model, Nras-mutant tumors (WHN89) were derived from *Tyr-CRE-ER^T2^ p16^L/L^ LSL-Nras^Q61R/Q61R^ (TpN^61R/61R^)* mouse model(*44*). *TpN^61R/61R^* tumors were minced and implanted into male C57Bl/6 mice.

Tumor volume was measured using digital calipers and calculated by length x width^2^/2. Mice in all treatment groups were euthanized when the vehicle-treated group reached the humane endpoint (tumor volume 1500-2000mm^3^).

#### PD0325901 treatment

Patient-derived xenografts were minced and subcutaneously injected into 8-week-old male NSG mice. When tumors reached 100-150 mm^3^, mice were randomly assigned into experimental groups. PD0325901 was prepared in 0.2% Tween 80 + 0.5% hydroxypropyl-methyl cellulose + 3% DMSO in ddH2O and administered by oral gavage (1.5 or 5 mg/kg/day).

#### Fenofibrate combination

M93-047 cells were subcutaneously injected into 8-week-old male NSG mice. One week after tumor cell inoculation (tumor size ∼ 40 mm^3^), mice were randomly assigned into five experimental groups: vehicle, fenofibrate, or fenofibrate + DHA. To study the effect of therapy in established tumors, M93-047 cells or minced patient-derived xenografts were subcutaneously injected into 4-week-old male NSG mice. When tumors reached 100-200 mm^3^, mice were randomly assigned into experimental groups. Syngeneic *TpN^61R/61R^* tumor cells (WHN89) were subcutaneously injected into 4-week-old male C57Bl/6 mice. When tumors reached ∼85 mm^3^, mice were randomly assigned into experimental groups and treated with vehicle, fenofibrate, DHA, or fenofibrate + DHA. Fenofibrate was prepared in 4% DMSO + 40% PEG300 + 2% Tween 80 in ddH2O and administered by oral gavage (200 mg/kg/day). DHA was administered by oral gavage (300 mg/kg/day) in soybean oil. Mice bearing patient-derived xenografts were given control chow, or DHA (0.15% w/w) plus FNB (0.15% w/w) laced chow (BioServ).

### Antibodies

#### Primary antibodies

Actin (Sigma-Aldrich #A5441 1:10000), COX-2 (PTGS2) (Santa Cruz Biotechnology, Inc. #sc-19999 1:200. FLAG (Sigma-Aldrich #F4049, 1 μg/mL/1:1000), HA (Sigma-Aldrich #H9658, flow cytometry 1:2000), 4-HNE (R&D Systems #MAB3249, flow cytometry 1:100, IF 10 μg/mL/1:100, IHC 20 μg/mL/1:50), NCoR1 (Bethyl #A301-146A, WB 1:1000; Sigma-Aldrich #PLA0170; Bethyl #A301-145A, PLA 10 mg/mL), 8-OHdG (Santa Cruz Biotechnology, Inc. #sc-66036, flow cytometry 1:100, IF 1:100, IHC 1:200), PPARα (Santa Cruz Biotechnology, Inc. #sc-398394, WB 1:500, PLA 1:100), S6K2 (Abcam #ab76960, WB 1:500; Abnova #H00006199-B02P, PLA 2.5 μg/mL), pIRE1α (Novus #2323, 1:2000). All other antibodies were purchased from Cell Signaling Technology. Dilution 1:1000, unless otherwise specified. pAkt (T308; #4056), BrdU (#5292, flow cytometry 1:200), pERK (T202/Y204; #4370), pMEK (S217/221; #9121), pRSK (T359/S363; #9344), S6 (#2317), pS6 (S235/236; #4858), pS6 (S240/244; #5364, WB 1:500, flow cytometry 1:100), S6K1 (#2708), pS6K1 (T389; #9234), S6K2 (#14130 WB 1:1000, PLA 1:50), XBP1s (#12782), BIP (#3177 WB), p-eIF2α (#3597 WB), eIF2α (#2103), ATF4 (#11815), CHOP (#2895, 1:500), PERK (#3192).

#### Secondary antibodies

WB: 1:10000. IRDye 680 Conjugated Goat (polyclonal) Anti-Rabbit IgG (H+L) (LI-COR #926-32211). IRDye 800CW Conjugated Goat (polyclonal) Anti-Mouse IgG (H+L) (LI-COR #926-32220). Flow cytometry/IF secondary antibodies were purchased from Life Technologies; 1:500. Alexa Fluor 488 goat anti-rabbit IgG (H+L) (#A11008). Alexa Fluor 594 goat anti-rabbit IgG (H+L) (#A11012). Alexa Fluor 647 goat anti-rabbit IgG (H+L) (#A21244). Alexa Fluor 488 goat anti-mouse IgG (H+L) (#A11029). Alexa Fluor 594 goat anti-rabbit IgG (H+L) (#A11032).

### Plasmids and lentivirus

Packaging plasmids pS-PAX2 and pMD2.G (a gift from Didier Trono (Addgene plasmid #12260, #12259)) were used to transfect 293T cells and produce lentiviral particles. Melanoma cells were transduced with lentivirus and selected with the appropriate antibiotics as previously described(61).

Plasmids: HA-tagged S6K1^T389EΔCT^ in pSLIK inducible vector was a gift from Kevin James (G418 selection, 2mg/mL; Addgene #58516). Protein expression was induced with doxycycline (0.25 μg/mL). Flag-tagged S6K1^T389S^ in pLJM vector was a gift from David Sabatini (puromycin selection, 2 μg/mL; Addgene #48800). HA-tagged S6K2^T388E^ (Addgene#17731) was subcloned into pLUT-poly inducible vector (puromycin selection, 2 μg/mL; gift from Dr. Meenhard Herlyn, The Wistar Institute). S6K1 and S6K2 short hairpin RNAs in pLKO.1 vector. S6K1 shRNA (sh1: TRCN0000003158, sh4: TRNC0000003161, sh5: TRNC0000003162). S6K2 shRNA (sh1: TRNC0000000729, sh4: TRNC0000010540, sh5: TRCN0000010541, sh6: TRCN0000199513, sh7: TRCN 0000432273).

### Immunoblotting

Cells were lysed with RIPA buffer (50 mM Tris-HCl, pH 7.4, 150 mM NaCl, 1% NP-40, 0.5% sodium deoxycholate, 1 mM EDTA, 2.5 mM sodium pyrophosphate, 0.05% or 0.1% SDS) supplemented with sodium vanadate (0.2 mM) and protease inhibitor cocktail (Sigma-Aldrich #11697498001). Immunoblotting was performed as previously described(45). Immunoblots were imaged with an Odyssey Infrared Imaging System (LI-COR Biosciences).

### Immunostaining

#### Flow cytometry

Cells were fixed in 90% ice cold EtOH/PBS and stored at -20°C. Cells were washed twice with PBS followed by 1 h incubation at RT with incubation buffer (1% BSA/PBS) containing primary antibodies (1:500 pS6 S240/244 Cell Signaling Technology #5364; 1:1000 HA Sigma-Aldrich #H9658). Samples were washed once (1 x PBS), incubated for 30 min at RT with incubation buffer containing secondary antibodies diluted to 1:500 (goat anti-mouse Alexa Fluor 488 ThermoFisher #A11029; goat anti-rabbit Alexa Fluor 647 #A21244), washed (1 x PBS) and resuspended in incubation buffer for analysis by flow cytometry (BD LSRII 14-color flow cytometer analyser; Alexa Fluor 488 and APC filter sets). Data analyses were performed using FlowJo software to calculate pS6 positivity. For 4-HNE staining, cells were fixed with 4% paraformaldehyde for 15 min, washed with PBS and stored at 4°C. Before immunostaining, cells were permeabilized with 0.5% Triton X-100/PBS for 10 min.

#### Immunohistochemistry

Tumors were fixed with 10% buffered formalin (MedSupply Partners #8BUFF-FORM 10%), embedded in paraffin and cut into 4 mm consecutive sections. Deparaffinized sections were steamed in citrate buffer (pH 6) for 20 min or incubated in pressure cooker with DAKO EDTA (pH9, Agilent Dako #S2367) for 20 min to retrieve antigens. Samples were treated with 3% H_2_O_2_ solution for 10 min and then Triton X-100 (0.3% in PBS) for 15 min at RT. Samples were blocked with 5% horse serum in PBS for 1h at RT, incubated with primary antibodies (4-HNE 1:50, 8-OHdG 1:200 in 4% BSA/PBS) overnight at 4°C, and incubated with secondary antibodies for 30 min at RT. All wash steps were performed with 0.5% PBST. Antigens were detected using diaminobenzidine (DAB). Slides were counterstained with hematoxylin. IHC images were acquired using a Nikon 80i upright microscope.

### Proximity ligation assay

Cells were grown on chamber slides (Lab-Tek II #154534), fixed with 4% paraformaldehyde for 10 min and washed three times with PBS. Proximity ligation assay was performed using Duolink In Situ Red Starter Kit Mouse/Rabbit (Sigma-Aldrich #DUO92101) per the manufacturer’s instructions. Slides were mounted and analyzed on a Leica TCS SP5 II scanning spectral confocal microscope.

### qRT-PCR

Cell pellets were homogenized using QIAshredder (QIagen #79656) and RNA extracted with PureLink RNA mini kit (Thermo Fisher Scientific, #12183018A). DNA was removed by on-column PureLink DNase treatment (Thermo Fisher Scientific, #12185010). First-strand cDNA was synthesized using Maxima cDNA kit (Thermo Scientific™ #K1642). Real time qPCR was performed using SYBR green PCR Master Mix (Applied Biosystems #4385612). Tumor samples for qRT-PCR were preserved in RNAlater solution (Invitrogen #AM7020). Primer sequences are specified in Supplemental Table 6.

### Cell viability assays

#### MTT assay

Cells were seeded in 96-well plates (2500 cells/well) and treated with drugs by adding 2X compounds (prepared in fresh growth medium) to the cells. After 72h, cell viability was determined by standard MTT Assays. Absorbance was determined at 490 nm using a BioTek plate reader.

#### CellTiter-Glo assay

Cells were seeded in 384-well plates (500 cells/well) and treated with drugs using the Janus MDT Nanohead. Viability was measured after 72h using CellTiter-Glo (Promega). Data were normalized where 100% viability equals luminescence of DMSO-treated cells, and 0% viability equals luminescence of 1 μM bortezomib-treated cells.

### Cell proliferation assay

#### Colorimetric detection

BrdU incorporation was performed per the manufacturer’s instructions (BrdU Cell Proliferation Assay Kit; Cell Signaling Technology #6813). Briefly, cells were seeded in 96-well plates (2500 cells/well) one day before drug treatment. Cells were treated with drugs for 72 h, followed by BrdU labelling for 16h and 30 min incubation in fixing/denaturing solution at RT. Cells were then incubated with primary antibody for 1h at RT, washed three times followed by incubation with HRP-conjugated antibody solution at RT for 30 min. Colorimetric reactions for BrdU incorporation were started by incubation with TMB substrate (10 min at RT) and terminated by adding STOP solution; signal was determined by absorbance at 450 nm.

### Cell death assay

Floating and attached cells were harvested, washed and stained with PSVue 643/PI or Annexin V-FITC/PI. Annexin V staining was performed as described previously (*45*), with 15 min Annexin V incubation. For PSVue/PI staining, cells were incubated in 3 μM PSVue 643/TES for 10 min at RT, washed in TES buffer and resuspended in PI (5 μg/mL)/TES for 15 min at RT. Cell death (Annexin V^+^/PSVue^+^ or PI^+^) was quantified by FACS (LSRII 18 color analyzer, BD Biosciences) and analyzed using FlowJo software v.10.7.2.

### Measurement of ROS production

Cells were trypsinized, washed with PBS and incubated with H2DCFDA [2’,7’-dichlorodihydrofluorescein diacetate, Thermo Fisher Scientific] (10 μM in PBS) for 25 min at 37°C. Cells were then washed once with PBS to remove the dye and incubated in phenol red-free growth medium for 10 min at 37°C. Following incubation, cells were placed on ice, protected from light and immediately analyzed by flow cytometry.

### Lipid peroxidation measurement

Cells were harvested and incubated with Bodipy 581/591 C11 (5 μM in growth medium; Thermo Fisher Scientific) for 30 min at 37°C. Cells were washed 5 times with PBS and then analyzed by flow cytometry.

### Proteomics

#### Sample preparation and LC-MS/MS analysis

M93-047 cells were transduced with vector control, S6K1 shRNA (sh1/4 or sh5) or S6K2 shRNA (sh1 or sh4) and lysed four days post-transduction with SDS buffer (50 mM Tris-HCl pH 7.5, 150 mM NaCl, 1% SDS and 1 mM EDTA) supplemented with protease inhibitors (150 mM PMSF, 1 mg/mL Pepstatin A and 1 mg/mL Leupeptin). In-gel trypsin digestion was performed using 15 mg(*62*).

LC-MS/MS of peptides was performed using a Waters nanoACQUITY UPLC in-line with a Thermo Q Exactive Plus mass spectrometer. For each sample, 1 mg of tryptic digest was loaded onto a 180 mm x 2 cm nanoACQUITY UPLC Symmetry C18 trap column with 5 mm particle size (Waters #186006527) followed by analytical separation on a 1.7 mm x 2.5 cm ACQUITY UPLC Peptide BEH C18 column with 1.7 µm particle size (Waters #186003546). Samples were analyzed in data-dependent mode using a 245 min LC gradient with water (solvent A) and acetonitrile (solvent B) containing 0.1% formic acid: 5-30% B over 225 min, 30-80% B over 5 min, 80% B hold for 10 min and return to initial conditions for 5 min. Full MS spectra were recorded at a resolution of 70,000 using a 400-2000 m/z scan range. Data-dependent MS/MS was performed on the top 20 most abundant ions selected with an isolation width of 1.5 m/z and were recorded at a resolution of 17,500.

#### Data analysis

RAW files were processed using MaxQuant 1.5.1.2 in a single run(*63*). Database searches were performed against the UniProt human sequence database (June 29, 2017; 159,819 sequences) and an in-house database of common laboratory contaminants (July 28, 2014; 3,671 sequences) with Trypsin/P specificity, a maximum of 2 missed cleavages and a minimum peptide length of 7 residues. Precursor and fragment mass tolerances were set to 4.5 ppm and 20 ppm, respectively. Protein N-terminal acetylation and methionine oxidation were set as variable modifications. Cysteine carbamidomethylation was set as a fixed modification. A maximum of 5 modifications were allowed per peptide. Peptide and protein false discovery rates were both set to 1%. Match between runs was enabled with a 0.7 min match time window and 20 min alignment time window. Protein quantification was based on unique peptides only. Label-free protein quantitation (LFQ) to normalize between samples was performed with a minimum peptide ratio of 1. Statistical analysis was performed using Perseus 1.5.0.31(*64*). Protein groups identified by a single unique peptide or corresponding to contaminants were removed. LFQ intensities were log2 transformed to reduce outlier effects, and missing values were imputed from a downshifted normal distribution. Statistical significance between conditions after pooling data from the two different shRNAs for the same S6K was defined as Student’s t-test p-value less than 0.05 and absolute fold-change greater than 2 based on log2-transformed LFQ intensities. Where indicated, correction for multiple testing was performed by permutation-based false discovery rate (FDR). Canonical pathway analysis was performed using QIAGEN’s Ingenuity® Pathway Analysis software (IPA®, QIAGEN Redwood City, www.qiagen.com/ingenuity) with all identified proteins as the reference dataset.

### Lipidomics

#### Sample preparation and LC-MS/MS analysis

M93-047 cells were transduced with vector control, S6K1 shRNA or S6K2 shRNA. Four days post transduction, lipids were extracted from cells grown in 10 cm plates using a modified Folch method(*65*). Briefly, media was removed from the plates and the cells were washed twice with PBS. Cells were quenched with ice-cold MeOH and transferred to a glass tube. Splash (Avanti Polar Lipids #330707) internal standard was added to each sample. Ice-cold 0.88% NaCl in water and CHCl_3_ were added to a final ratio of 2:1:1 CHCl_3_:MeOH:0.88% NaCl. Extracts were then sonicated and centrifuged. The lower organic phase was transferred to a new glass tube. The upper aqueous phase was re-extracted with synthetic organic phase, and the re-extracted organic phase was pooled with the initial extract. The pooled organic phases were dried under nitrogen, and lipids were reconstituted in 1:8:1 CHCl_3_:MeOH:H_2_O. Global lipid extracts from triplicate plates were analyzed by reversed-phase LC-MS/MS on a Thermo Q Exactive HF-X in both positive and negative polarities. For each polarity, 1% (2 μL) of the resuspended lipidome was loaded onto a 150 μm x 2.1 cm Accucore C30 column with a 2.6 μm particle size (Thermo Fisher Scientific #27826-152130). The lipidome was analyzed using data-dependent acquisition on a 40-minute gradient with 1:1 acetonitrile:water with 5 mM ammonium formate and 0.1% formic acid (solvent A) and 88:10:2 isopropanol:acetonitrile:water with 5 mM ammonium formate and 0.1% formic acid (solvent B). The gradient is as follows: 0-60% B over 10 minutes, 60-85% B over 10 minutes, 85-100% over 10 minutes and holding for 5 minutes and re-equilibrating to initial conditions over 5 minutes. Full MS spectra were recorded at a resolution of 120,000 using a 300-1,200 m/z scan range in positive mode and 250-1,200 m/z scan range in negative mode. Data-dependent MS/MS scans were recorded at a resolution of 15,000 for the top 20 most abundant ions isolated with a 0.4 m/z isolation width. Stepped collision energy (NCE) of 20/30 was used in positive mode and 20/30/40 was used in negative mode.

#### Data analysis

Quantitation for global lipids was performed using Lipid Search 4.1 (Thermo Fisher Scientific). Briefly, raw files were processed using Lipid Search software in a single batch. Lipids were identified based on mass, retention time and predicted fragmentation patterns. All identified lipids are aligned and quantified with a 5 ppm mass tolerance and a retention time window of 0.2 minutes. Manual filtering included removing low scoring lipid identifications and unlikely adducts based on standards. To quantify lipid distribution, the abundance (peak area) of each lipid species in a sample was normalized to total lipid peak area in that sample. Statistical significance was defined as a Student’s t-test p-value less than 0.05.

### RNA seq

#### Sample preparation

Cells were lysed in Tri-Reagent (Sigma-Aldrich, #T9424) and RNA extracted using the Direct-zol MiniPrep Kit (Zymo Research, #R2050) with on-column DNAse I treatment. Total RNA (200ng) was used to prepare 3’ QuantSeq mRNASeq libraries (Lexogen, #KO152×96) as per protocol. Fourteen PCR cycles were used for amplification during the library preparation. Libraries were checked for quality using the TapeStation High Sensitivity D5000 ScreenTape (Agilent Technologies, #5067-5592). Quantitation of library was done using qPCR (Roche #KK4835). Sequencing was done on the NextSeq 500 (Illumina) per manufacturer’s instructions using a 75 cycle High Output sequencing kit. A final concentration of 1.9pM was loaded onto the flowcell.

#### Data analysis

RNA-seq data was aligned using *bowtie2*(*66*) against hg19 version of the human genome and *RSEM* v1.2.12 software(*67*) was used to estimate raw read counts and RPKM using Ensemble transcriptome. *DESeq2*(*68*) was used to estimate significance of differential expression between each shRNA and control condition. Genes with expression changes passing FDR<5% threshold were considered significant. Only genes that overlapped between the two shRNAs for S6K1 (sh1/4 and sh5) or S6K2 (sh1 and sh4) were analyzed. Heatmaps were normalized by DESeq2 log2 scaled counts relative to average across all samples. Gene set enrichment analysis was done using QIAGEN’s Ingenuity® Pathway Analysis software and significance of enrichment for Upstream Regulators was defined at p-value<0.05.

### Reverse phase protein array

Cells were lysed with 1% Triton X-100, 50 mM HEPES, pH 7.4, 150 mM NaCl, 1.5 mM MgCl_2_, 1 mM EGTA, 10% glycerol, protease and phosphatase cocktail (Sigma-Aldrich #11873580001 and 04906837001). Lysates were denatured in sample buffer (10% glycerol, 2% SDS, 0.06 M Tris-HCl, pH 6.8, 1/40th volume of 2-ME) for 5 min, 95°C. RPPA was performed by the MD Anderson Center RPPA core facility as previously described(*69*).

### TCGA data, S6K1 and S6K2 correlation and survival

To determine the relationship between S6K2 and S6K1 expression in human melanoma patient tumors, log2-transformed mRNA expression values for S6K1 and S6K2 were extracted from the TCGA Skin Cutaneous Melanoma (SKCM) dataset (n=443) via cBioPortal (accessed 5^th^ Feb 2022).

Kaplan–Meier survival curves for high vs. low S6K1 or S6K2 expression in melanoma patients (n=470) were obtained using the TCGA skin cutaneous melanoma dataset. The analysis was performed using R2: Genomics Analysis and Visualization Platform (http://r2.amc.nl) to stratify by first (low, n=115) vs. last (high, n=115) quartiles. P values correspond to log-rank tests comparing the two Kaplan–Meier curves.

### Tumor growth rate Analysis

For the analysis of longitudinal data from *in vivo* tumor growth experiments, linear mixed effects models were applied to compare tumor growth rates between treatment groups. For multiple comparisons, FDR adjusted P-value <0.05 was considered significant.

## QUANTIFICATION AND STATISTICAL ANALYSIS

Data were analyzed and plotted using GraphPad Prism or Microsoft Excel. Data are presented as mean ± SD or median with interquartile range as indicated in each figure legend; replicate numbers (n) are indicated in the figure legends. In graphs showing mean of multiple independent experiments, values of technical replicates were first averaged to represent each experiment, and ‘n’ denotes how many independent experiments were performed. In graphs showing one representative experiment, ‘n’ denotes number of biological replicates. Statistical differences between two experimental groups were calculated using the unpaired two-tailed Student’s t-test, unless otherwise specified.

## DATA AND SOFTWARE AVAILABILITY

Proteomics dataset generated from M93-047 cell line has been deposited to the MassIVE repository (accession # MSV000083480). https://massive.ucsd.edu/ProteoSAFe/dataset.jsp?task=e702a8018b224a52a737ee2ba323b3a4

RNA-seq dataset generated from M93-047 cell line has been deposited into the Gene Expression Omnibus (GEO) repository (accession #GSE127916). https://www.ncbi.nlm.nih.gov/geo/query/acc.cgi?acc=GSE127916

## Supplementary figure legends

**Fig. S1. Related to Fig.1.**
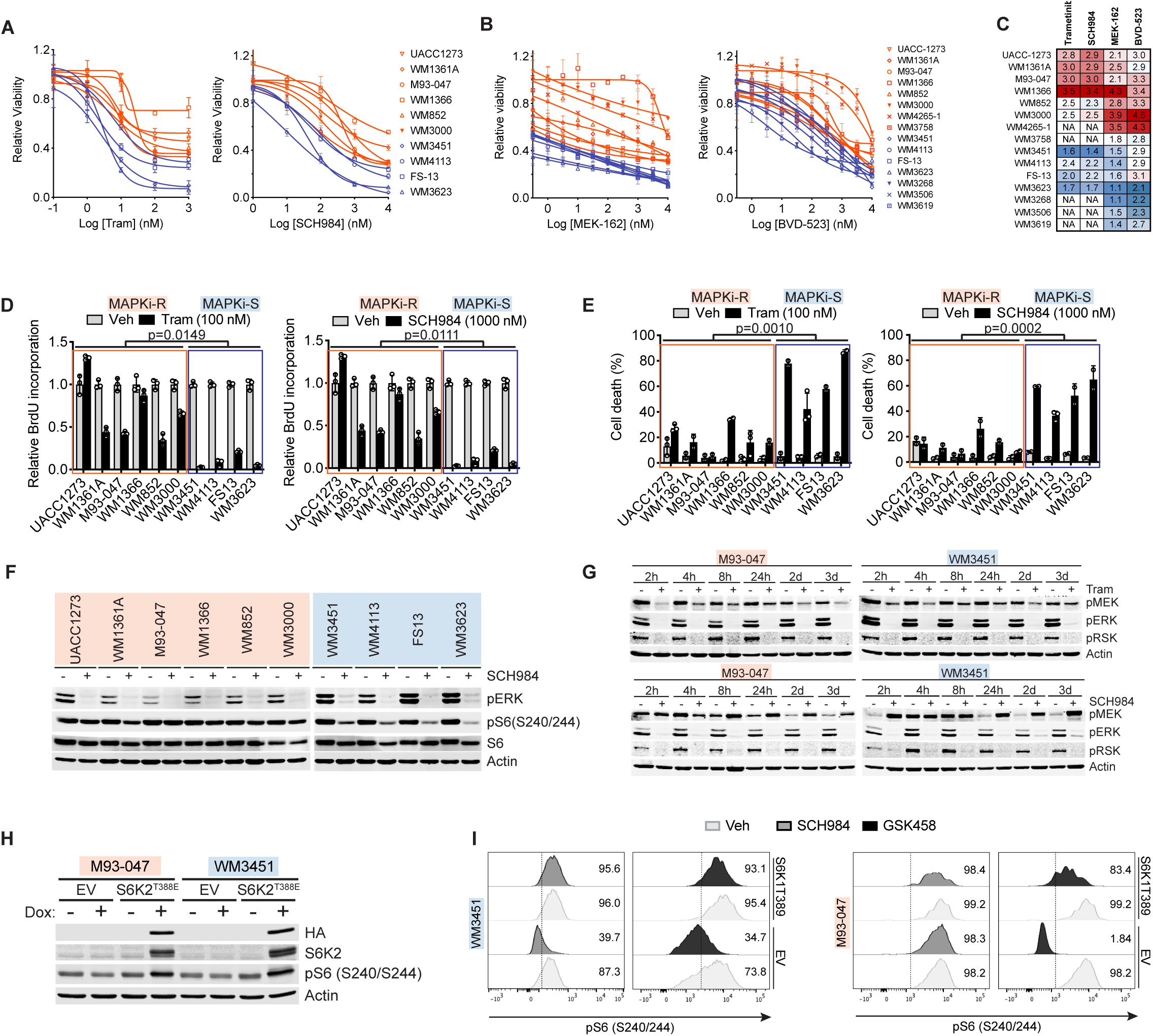
**A.-C.** NRAS-mutant melanoma cells were treated with MAPK inhibitors to determine dose-response relationships. Cell viability was determined 72h post-treatment by (**A**) MTT (trametinib; SCH772984) (mean ± SD; n=7) or (**B**) CellTiter-Glo assays (MEK-162; BVD-523) (mean ± SD; n=2). (**C**) Sensitivity of NRAS-mutant melanoma cells to MAPK inhibitors (**A, B**) represented as area under the dose-response curve values in a blue-to-red heatmap with white median. **D., E.** MAPKi-S or MAPKi-R cells were treated with trametinib (100 nM) or SCH772984 (1000 nM) for 72h. (**D**) BrdU incorporation relative to DMSO; mean ± SD (n=3). The difference in BrdU incorporation between DMSO-and drug-treated cells was calculated for each cell line. MAPKi-S *vs.* MAPKi-R compared by unpaired, two-tailed Student’s *t*-test. (**E**) Cell death was assessed by PSVue/PI staining (n=2). Cell death induction was calculated in each cell line and compared in MAPKi-R *vs.* MAPKi-S cells by unpaired, two-tailed Student’s *t*-test. **F.** MAPKi-R or MAPKi-S cells were treated with SCH772984 (1000 nM) for 24h and analyzed by immunoblotting. **G.** Kinetics of MAPK pathway inhibition. Cells treated trametinib (100 nM) or SCH772984 (1000 nM) were analyzed by immunoblotting. **H.** Cells were transduced with doxycycline-inducible S6K2^T388E^ and incubated with 0.25 µg/ml doxycycline for 48h. S6K2 and phospho-S6 (S240/244) levels were determined by immunoblotting **i** Cells transduced with doxycycline-inducible S6K1T389E were incubated with doxycycline for 24h and then treated with ERKi SCH984 (1000 nM) or PI3K/mTORi GSK458 (100 nM) for an additional 24h. Percentage of pS6(^S240/244^)+ cells was assessed by flow cytometry (n=1).

**Fig. S2. Related to Fig.3.**
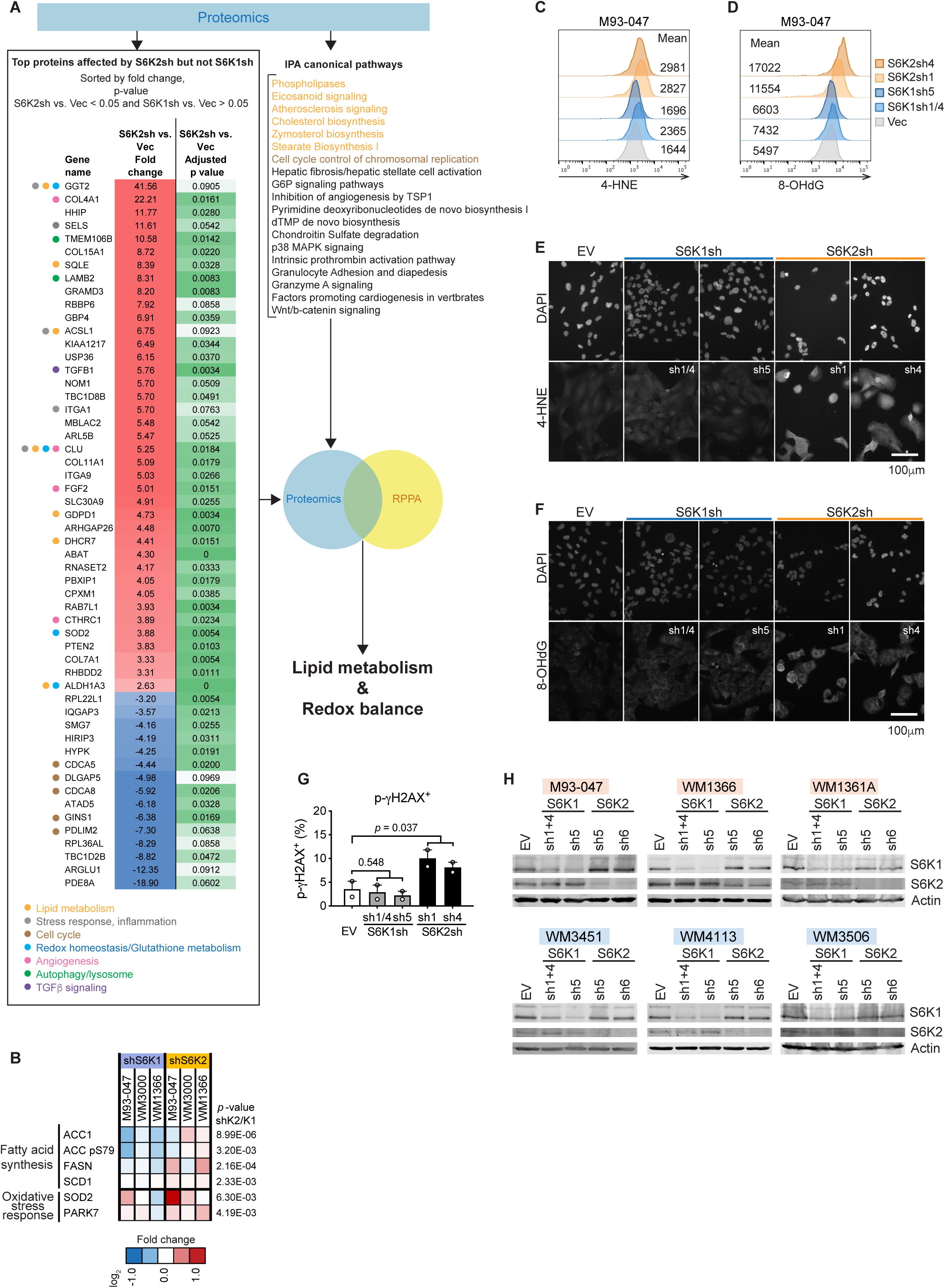
**A.** NRAS-mutant melanoma cells (M93-047) were transduced with lentivirus encoding S6K1 shRNA, S6K2 shRNA or non-targeting vector control (Vec) and subjected to LC-MS/MS (**A**) or reversed phase protein array (RPPA) (**B**) analysis (4 dpi). **A.** Schematic depicts the highest ranked proteins differentially regulated by S6K2 *vs.* S6K1 knockdown, identifying lipid metabolism and redox homeostasis as processes disrupted by S6K2 depletion. Pathways/biological processes for the top proteins were overlaid with IPA canonical pathway analysis, results are color coded. **B.** Heatmap depicts log_2_ mean fold change of 2 shRNA relative to vector for lipid metabolism and oxidative stress response targets in the RPPA dataset. Significant differences between S6K1sh vs. S6K2sh were estimated using paired, two-sided t-tests. **C.-G.** M93-047 cells were transduced with lentivirus encoding S6K1 or S6K2 shRNA and stained for the lipid peroxidation product 4-hydroxynonenal (4-HNE) (**C, E**) or the oxidative DNA damage marker 8-OHdG (**D, F**) 6 dpi. Samples were analyzed by flow cytometry (**C-D;** mean fluorescence intensity is shown) or immunofluorescence (**E-F**). **G.** DNA double strand breaks were assessed by p-gH2AX flow cytometry 6 dpi (mean ± SD, n=2). **H.** Representative Western blots (n=2) showing knockdown of S6K1 or S6K2 (4 dpi) for **Fig. 3 C-F** samples. **A, B, G.** Statistical significance was assessed by unpaired two-tailed Student’s *t*-tests. A 10% FDR was applied and Benjamini-Hochberg adjusted p values reported (**a**).

**Fig. S3. Related to Fig.4.**
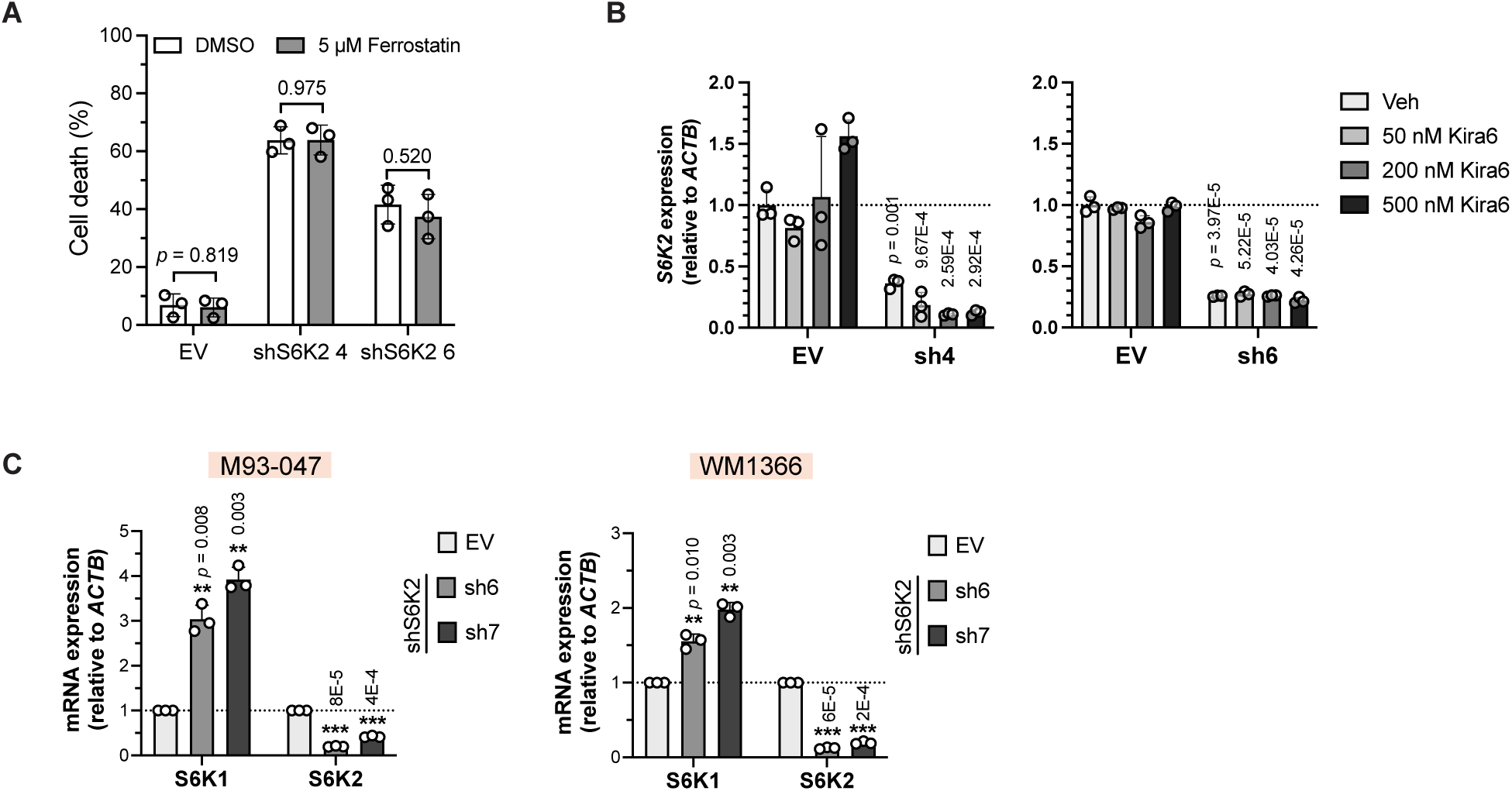
**A.** M93-047 cells were treated as in Fig. **4D**, except media was supplemented with ferrostatin to evaluate ferroptosis dependence. **B.** S6K2 levels in samples from Fig. **4C** were monitored by qRT-PCR. **C.** qRT-PCR analysis of S6K1 and S6K2 levels from Fig. **4F**. **A, B, C**, Data are mean ± SD from three independent experiments. *P*-values from unpaired two-sided *t*-tests.

**Fig. S4. Related to Fig.5.**
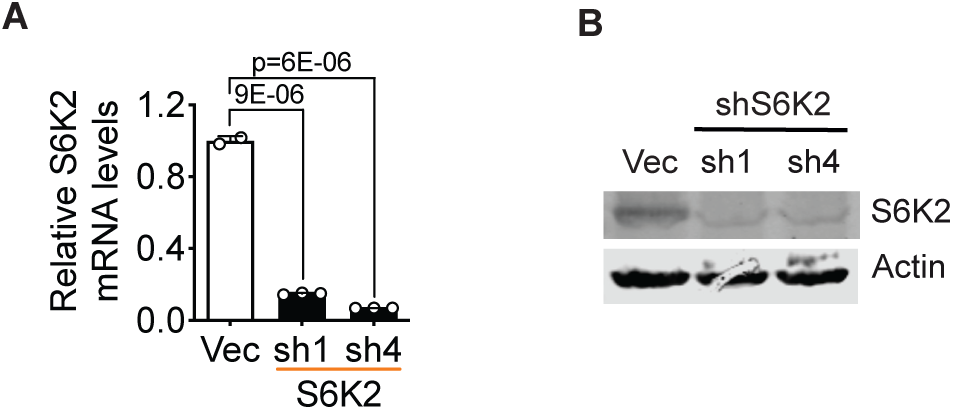
**A.** S6K2 knockdown efficiency was assessed by by qRT-PCR (Fig 5**D**; 4dpi); mean ± SD (n=2-3) **B.** S6K2 knockdown efficiency of Fig 5**E** (4dpi) was assessed by western blotting.

**Fig. S5. Related to Fig.6.**
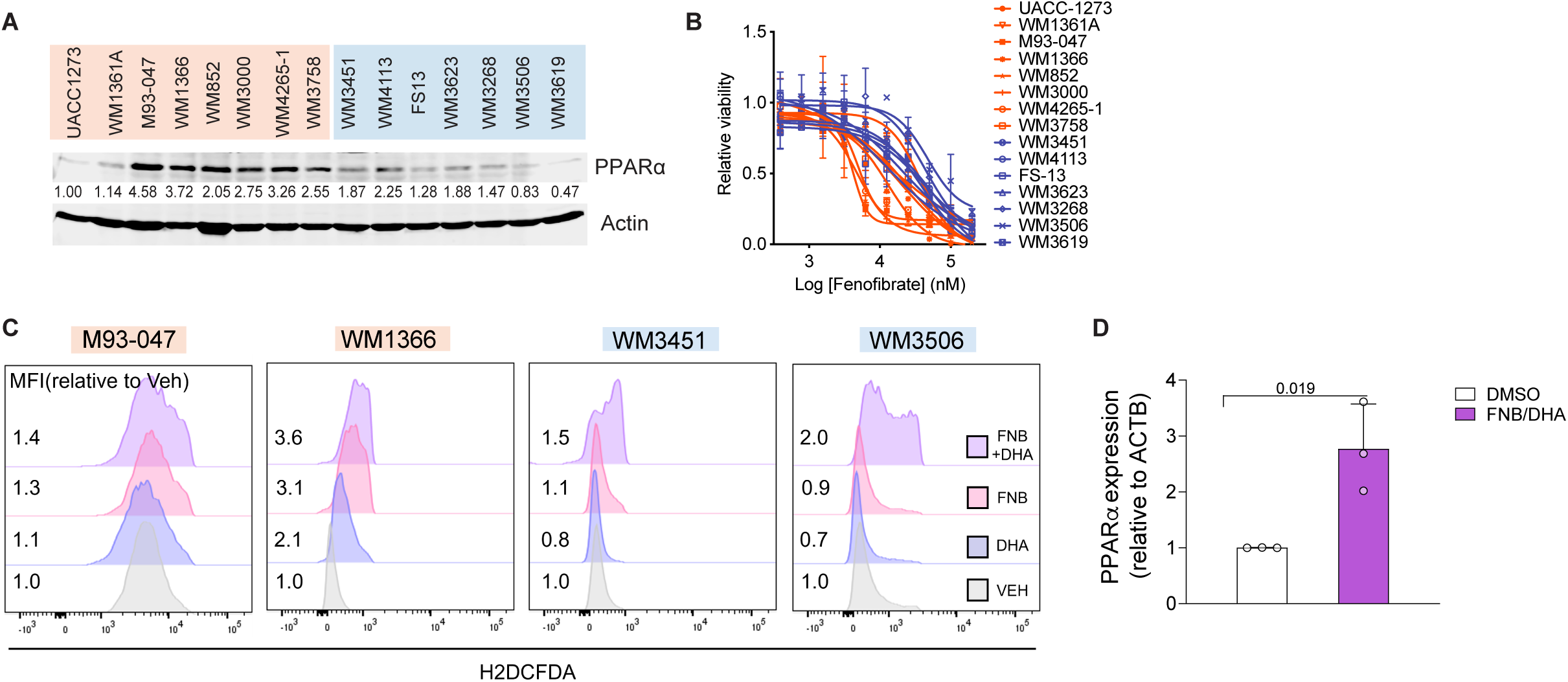
**A.** PPARα levels in MAPKi-R and MAPKi-S cells were determined by western blotting. PPARα levels were normalized to actin; numbers indicate protein levels relative to UACC1273. **B.** MAPKi-R (red) and MAPKi-S (blue) cells were treated with increasing doses of fenofibrate for 72h; cell viability was determined by CellTiter-Glo. Data represent average from two independent experiments ± SD. **C.** Cells were treated with 50 µM fenofibrate, 7.5 µM DHA as single agents or in combination. ROS was assessed by H2DCFDA. Representative histograms (n=3) showing mean fluorescence intensity relative to vehicle control. **D.** Levels of PPARα were quantified by qRT-PCR mean SD (n=3); unpaired two-sided *t* test

**Fig. S6 Related to Fig.7.**
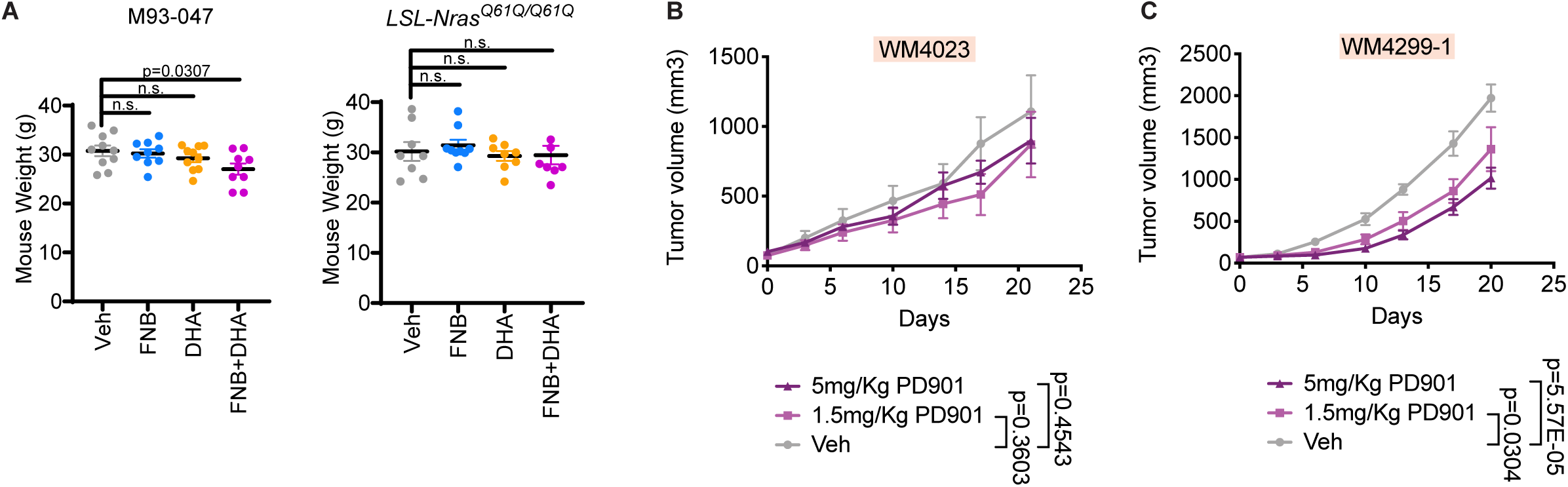
**A.** Weight of animals enrolled in the study depicted in Fig 7a,b at the last follow-up time points (day 24 and 30 respectively). Mice were treated with fenofibrate (200mg/kg) or DHA (300mg/kg) as a single agent or in combination. Data are shown as mean ± SEM**. B., C.** Mice bearing WM4023 or WM4229-1 patient derived xenografts (∼85 or 70 mm^3^) were treated with the MEK inhibitor (PD0325901 (1.5 or 5 mg/kg) and classified as MAPKi-resistant PDXs; mean ± SEM, n=4 mice/treatment group. FDR adjusted p-values were estimated from a linear mixed-effect model with all follow-up subjects and time points.

## REFERENCES

1. N. K. Hayward et al., Whole-genome landscapes of major melanoma subtypes.Nature 545, 175–180 (2017).

2. R. Dummer et al., Binimetinib versus dacarbazine in patients with advanced NRAS-mutant melanoma (NEMO): a multicentre, open-label, randomised, phase 3 trial. Lancet Oncol 18, 435–445 (2017).

3. R. J. Sullivan et al., First-in-Class ERK1/2 Inhibitor Ulixertinib (BVD-523) in Patients with MAPK Mutant Advanced Solid Tumors: Results of a Phase I Dose-Escalation and Expansion Study. Cancer Discov 8, 184–195 (2018).

4. H. L. Vu, A. E. Aplin, Targeting mutant NRAS signaling pathways in melanoma. Pharmacol Res 107, 111–116 (2016).

5. <Activation of an early feedback survival loop involving phospho ErbB3.pdf>.

6. Y. N. Gopal et al., Basal and treatment-induced activation of AKT mediates resistance to cell death by AZD6244 (ARRY-142886) in Braf-mutant human cutaneous melanoma cells. Cancer Res 70, 8736–8747 (2010).

7. M. Atefi et al., Reversing melanoma cross-resistance to BRAF and MEK inhibitors by co-targeting the AKT/mTOR pathway. PLoS One 6, e28973 (2011).

8. D. Juric et al., A phase 1b dose-escalation study of BYL719 plus binimetinib (MEK162) in patients with selected advanced solid tumors. 32, 9051–9051 (2014).

9. A. W. Tolcher et al., Phase I study of the MEK inhibitor trametinib in combination with the AKT inhibitor afuresertib in patients with solid tumors and multiple myeloma. Cancer Chemother Pharmacol 75, 183–189 (2015).

10. C. Posch et al., Searching for the Chokehold of NRAS Mutant Melanoma. J Invest Dermatol 136, 1330–1336 (2016).

11. A. Karz et al., Melanoma central nervous system metastases: An update to approaches, challenges, and opportunities. Pigment Cell Melanoma Res 35, 554–572 (2022).

12. A. P. Algazi et al., Dual MEK/AKT inhibition with trametinib and GSK2141795 does not yield clinical benefit in metastatic NRAS-mutant and wild-type melanoma. Pigment Cell Melanoma Res 31, 110–114 (2018).

13. C. Posch et al., Combined targeting of MEK and PI3K/mTOR effector pathways is necessary to effectively inhibit NRAS mutant melanoma in vitro and in vivo. Proc Natl Acad Sci U S A 110, 4015–4020 (2013).

14. K. Duvel et al., Activation of a metabolic gene regulatory network downstream of mTOR complex 1. Mol Cell 39, 171–183 (2010).

15. R. B. Corcoran et al., TORC1 suppression predicts responsiveness to RAF and MEK inhibition in BRAF-mutant melanoma. Sci Transl Med 5, 196ra198 (2013).

16. B. Magnuson, B. Ekim, D. C. Fingar, Regulation and function of ribosomal protein S6 kinase (S6K) within mTOR signalling networks. Biochem J 441, 1–21 (2012).

17. S. Sridharan, A. Basu, Distinct Roles of mTOR Targets S6K1 and S6K2 in Breast Cancer. Int J Mol Sci 21, (2020).

18. O. E. Pardo, M. J. Seckl, S6K2: The Neglected S6 Kinase Family Member. Front Oncol 3, 191 (2013).

19. X. Chen et al., Integrative proteomic and phosphoproteomic profiling of invasive micropapillary breast carcinoma. J Proteomics 257, 104511 (2022).

20. S. Sridharan, A. Basu, S6 kinase 2 promotes breast cancer cell survival via Akt. Cancer Res 71, 2590–2599 (2011).

21. M. Pende et al., S6K1(-/-)/S6K2(-/-) mice exhibit perinatal lethality and rapamycin-sensitive 5’-terminal oligopyrimidine mRNA translation and reveal a mitogen-activated protein kinase-dependent S6 kinase pathway. Mol Cell Biol 24, 3112–3124 (2004).

22. <Disruption of the p70s6k p85s6k gene reveals a small.pdf>.

23. I. C. Pavan et al., Different interactomes for p70-S6K1 and p54-S6K2 revealed by proteomic analysis. Proteomics 16, 2650–2666 (2016).

24. E. Karlsson et al., Revealing Different Roles of the mTOR-Targets S6K1 and S6K2 in Breast Cancer by Expression Profiling and Structural Analysis. PLoS One 10, e0145013 (2015).

25. C. Nardella et al., Differential expression of S6K2 dictates tissue-specific requirement for S6K1 in mediating aberrant mTORC1 signaling and tumorigenesis. Cancer Res 71, 3669–3675 (2011).

26. P. A. Ascierto et al., MEK162 for patients with advanced melanoma harbouring NRAS or Val600 BRAF mutations: a non-randomised, open-label phase 2 study. Lancet Oncol 14, 249–256 (2013).

27. <Characterization of S6K2, a novel kinase homologous to S6K1.pdf>.

28. A. Nagler et al., A genome-wide CRISPR screen identifies FBXO42 involvement in resistance toward MEK inhibition in NRAS-mutant melanoma. Pigment Cell Melanoma Res 33, 334–344 (2020).

29. A. Ayala, M. F. Munoz, S. Arguelles, Lipid peroxidation: production, metabolism, and signaling mechanisms of malondialdehyde and 4-hydroxy-2-nonenal. Oxid Med Cell Longev 2014, 360438 (2014).

30. L. Magtanong, P. J. Ko, S. J. Dixon, Emerging roles for lipids in non-apoptotic cell death. Cell Death Differ 23, 1099–1109 (2016).

31. S. J. Dixon et al., Ferroptosis: an iron-dependent form of nonapoptotic cell death. Cell 149, 1060–1072 (2012).

32. W. S. Yang et al., Regulation of ferroptotic cancer cell death by GPX4. Cell 156, 317–331 (2014).

33. V. S. Viswanathan et al., Dependency of a therapy-resistant state of cancer cells on a lipid peroxidase pathway. Nature 547, 453–457 (2017).

34. K. Shimada et al., Global survey of cell death mechanisms reveals metabolic regulation of ferroptosis. Nat Chem Biol 12, 497–503 (2016).

35. J. P. Friedmann Angeli et al., Inactivation of the ferroptosis regulator Gpx4 triggers acute renal failure in mice. Nat Cell Biol 16, 1180–1191 (2014).

36. S. Kersten, Integrated physiology and systems biology of PPARalpha. Mol Metab 3, 354–371 (2014).

37. M. Rakhshandehroo, B. Knoch, M. Muller, S. Kersten, Peroxisome proliferator-activated receptor alpha target genes. PPAR Res 2010, (2010).

38. J. Han, R. J. Kaufman, The role of ER stress in lipid metabolism and lipotoxicity. J Lipid Res 57, 1329–1338 (2016).

39. A. Almanza et al., Endoplasmic reticulum stress signalling - from basic mechanisms to clinical applications. FEBS J 286, 241–278 (2019).

40. A. Sengupta et al., Targeted disruption of glutathione peroxidase 4 in mouse skin epithelial cells impairs postnatal hair follicle morphogenesis that is partially rescued through inhibition of COX-2. J Invest Dermatol 133, 1731–1741 (2013).

41. K. Kim, S. Pyo, S. H. Um, S6 kinase 2 deficiency enhances ketone body production and increases peroxisome proliferator-activated receptor alpha activity in the liver. Hepatology 55, 1727–1737 (2012).

42. <PPAR agonist fenofibrate suppresses tumor growth.pdf>.

43. X. Lian et al., Anticancer Properties of Fenofibrate: A Repurposing Use. J Cancer 9, 1527–1537 (2018).

44. C. E. Burd et al., Mutation-specific RAS oncogenicity explains NRAS codon 61 selection in melanoma. Cancer Discov 4, 1418–1429 (2014).

45. I. M. Echevarria-Vargas et al., Co-targeting BET and MEK as salvage therapy for MAPK and checkpoint inhibitor-resistant melanoma. EMBO Mol Med 10, (2018).

46. E. Szegezdi, S. E. Logue, A. M. Gorman, A. Samali, Mediators of endoplasmic reticulum stress-induced apoptosis. EMBO Rep 7, 880–885 (2006).

47. M. J. Hangauer et al., Drug-tolerant persister cancer cells are vulnerable to GPX4 inhibition. Nature 551, 247–250 (2017).

48. J. Tsoi et al., Multi-stage Differentiation Defines Melanoma Subtypes with Differential Vulnerability to Drug-Induced Iron-Dependent Oxidative Stress. Cancer Cell 33, 890–904 e895 (2018).

49. J. Tsoi et al., Multi-stage Differentiation Defines Melanoma Subtypes with Differential Vulnerability to Drug-Induced Iron-Dependent Oxidative Stress. Cancer Cell, (2018).

50. S. Gerstenecker et al., Discovery of a Potent and Highly Isoform-Selective Inhibitor of the Neglected Ribosomal Protein S6 Kinase Beta 2 (S6K2). Cancers (Basel) 13, (2021).

51. J. Villanueva et al., Concurrent MEK2 mutation and BRAF amplification confer resistance to BRAF and MEK inhibitors in melanoma. Cell Rep 4, 1090–1099 (2013).

52. J. L. F. Teh et al., In Vivo E2F Reporting Reveals Efficacious Schedules of MEK1/2-CDK4/6 Targeting and mTOR-S6 Resistance Mechanisms. Cancer Discov 8, 568–581 (2018).

53. G. Romano et al., A Preexisting Rare PIK3CA(E545K) Subpopulation Confers Clinical Resistance to MEK plus CDK4/6 Inhibition in NRAS Melanoma and Is Dependent on S6K1 Signaling. Cancer Discov 8, 556–567 (2018).

54. D. R. Green, The Coming Decade of Cell Death Research: Five Riddles. Cell 177, 1094–1107 (2019).

55. M. Cerezo et al., Compounds Triggering ER Stress Exert Anti-Melanoma Effects and Overcome BRAF Inhibitor Resistance. Cancer Cell 29, 805–819 (2016).

56. L. Wang et al., An Acquired Vulnerability of Drug-Resistant Melanoma with Therapeutic Potential. Cell 173, 1413–1425.e1414 (2018).

57. N. Rufo, A. D. Garg, P. Agostinis, The Unfolded Protein Response in Immunogenic Cell Death and Cancer Immunotherapy. Trends in Cancer 3, 643–658 (2017).

58. L. Liu et al., Ablation of ERO1A induces lethal endoplasmic reticulum stress responses and immunogenic cell death to activate anti-tumor immunity. Cell Rep Med 4, 101206 (2023).

59. J. Y. Lee et al., Polyunsaturated fatty acid biosynthesis pathway determines ferroptosis sensitivity in gastric cancer. Proc Natl Acad Sci U S A 117, 32433–32442 (2020).

60. S. Doll et al., ACSL4 dictates ferroptosis sensitivity by shaping cellular lipid composition. Nat Chem Biol 13, 91–98 (2017).

61. P. Reyes-Uribe et al., Exploiting TERT dependency as a therapeutic strategy for NRAS-mutant melanoma. Oncogene 37, 4058–4072 (2018).

62. L. A. Beer, H. Y. Tang, S. Sriswasdi, K. T. Barnhart, D. W. Speicher, Systematic discovery of ectopic pregnancy serum biomarkers using 3-D protein profiling coupled with label-free quantitation. J Proteome Res 10, 1126–1138 (2011).

63. J. Cox, M. Mann, MaxQuant enables high peptide identification rates, individualized p.p.b.-range mass accuracies and proteome-wide protein quantification. Nat Biotechnol 26, 1367–1372 (2008).

64. S. Tyanova et al., The Perseus computational platform for comprehensive analysis of (prote)omics data. Nat Methods 13, 731–740 (2016).

65. J. Folch, M. Lees, G. H. Sloane Stanley, A simple method for the isolation and purification of total lipides from animal tissues. J Biol Chem 226, 497–509 (1957).

66. B. Langmead, S. L. Salzberg, Fast gapped-read alignment with Bowtie 2. Nat Methods 9, 357–359 (2012).

67. B. Li, C. N. Dewey, RSEM: accurate transcript quantification from RNA-Seq data with or without a reference genome. BMC Bioinformatics 12, 323 (2011).

68. M. I. Love, W. Huber, S. Anders, Moderated estimation of fold change and dispersion for RNA-seq data with DESeq2. Genome Biol 15, 550 (2014).

69. R. Tibes et al., Reverse phase protein array: validation of a novel proteomic technology and utility for analysis of primary leukemia specimens and hematopoietic stem cells. Mol Cancer Ther 5, 2512–2521 (2006).

