## Supplementary Table 1 for "Selective abrogation of S6K2 maps lipid homeostasis as a survival vulnerability in MAPKi-resistant NRAS^MUT^ melanoma"

Table S1: Related to Fig. 1 and Fig. S1. List of NRAS-mutant human melanoma cell lines.

| **Cell line** | **NRAS** | | **BRAF** |
| --- | --- | --- | --- |
| M93-047 | Q61K | Het | WT |
| WM1366 | Q61L | Hom | WT |
| WM1361A | Q61R | Hom | WT |
| WM852 | Q61R | Hom | WT |
| WM4265 | Q61K | N/D | WT |
| WM3000 | Q61R | Het | WT |
| UACC1273 | Q61L | Het | WT |
| WM3758 | Q61L | Het | WT |
| WM3451 | Q61K | Het | WT |
| WM4113 | Q61R | Hom | WT |
| WM3506 | Q61R | N/D | WT |
| WM3268 | Q61K | Het | WT |
| WM3619 | Q61R | Hom | WT |
| WM3623 | Q61K | Het | WT |
| FS13 | Q61L | Het | WT |
