## Supplementary Table 5 for "Selective abrogation of S6K2 maps lipid homeostasis as a survival vulnerability in MAPKi-resistant NRAS^MUT^ melanoma"

**Supplementary Table 5. Related to Fig. 4a. List of overrepresented genes and z-scores predicting UPR activation and increased/inhibited function of PPARs in S6K2 or S6K1-depleted cells.**

|  | Type | Regulator | P | Z | N | pos | neg | Molecules |
| --- | --- | --- | --- | --- | --- | --- | --- | --- |
| S6K2sh | transcription regulator | XBP1 | 2.42E-02 | 4.068 | 20 | 19 | 1 | BECN1(+1.368),BET1(+2.560),BLZF1(+2.445),DNAJC1(+1.556),DNAJC3(+1.563),EIF2AK3(+1.709),ERLEC1(+1.575),FKBP14(+1.743),LMAN1(+1.372),MAP1LC3B(+1.405),SEC11A(+1.548),SEC63(+1.366),SELENOM(+1.805),SERP1(+1.648),SPARC(+1.447),SPCS3(+2.027),TOP1(+1.441),TRAM1(+2.059),XRCC6(-1.788),YIPF5(+1.342) |
|  | other | TXNIP | 2.40E-04 | 1.446 | 10 | 7 | 3 | BCL6(+1.817),CCNA2(-5.214),CCND1(-1.599),CDKN1B(+1.821),DDIT4(+1.559),IL3RA(+24.021),PCNA(-1.843),PTGS2(+3.339),STARD4(+1.501),VEGFA(+1.835) |
|  | kinase | PERK | 3.83E-03 | 2.05 | 14 | 11 | 3 | BECN1(+1.368),BTG2(+2.180),CCND1(-1.599),CLCN3(+1.625),DNAJC3(+1.563),EIF2AK3(+1.709),FGF2(+2.213),FOSL1(-1.405),MAP1LC3B(+1.405),MTHFR(+2.056),PON2(+1.406),RPS6KA2(+1.989),SHMT2(-1.429),VEGFA(+1.835) |
|  | transcription regulator | ATF4 | 9.68E-03 | 2.021 | 17 | 12 | 5 | ABCA1(+2.225),AREG(+2.001),BBC3(+1.564),CDKN1B(+1.821),CHAC1(+1.823),DDIT4(+1.559),LGALS3(+2.070),MAP1LC3B(+1.405),PMP22(+1.476),PRKDC(-1.638),PTGS2(+3.339),SHMT2(-1.429),SLC1A5(-1.985),SLC38A2(+1.913),SLC7A1(-1.544),SLC7A5(-3.485),VEGFA(+1.835) |
|  | chemical drug | pioglitazone | 1.79E-01 | 1.767 | 13 | 6 | 7 | *ABCA1*(+2.225),*BRCA1*(-2.980),*CCND1*(-1.599),*CDKN1B*(+1.821),*EPAS1*(+1.845),*FOSL1*(-1.405),*NCEH1*(+1.653),*NR4A3*(+3.464),*PCNA*(-1.843),*PTGS2*(+3.339),*SERPINE1*(-1.537),*STAT5B*(-1.637),*TNFRSF1B*(-2.495) |
|  | chemical drug | ciglitazone | 9.97E-02 | 1.833 | 8 | 4 | 4 | *ABCA1*(+2.225),*BCL2L1*(-1.951),*CCND1*(-1.599),*CDKN1B*(+1.821),*MMP2*(+1.598),*PARP1*(-1.794),*PTGS2*(+3.339),*SERPINE1*(-1.537) |
|  | chemical drug | troglitazone | 2.68E-04 | 1.276 | 40 | 10 | 30 | *ABCA1*(+2.225),*BCL2L1*(-1.951),*BIRC5*(-3.100),*C3*(+32.006),*CCNB1*(-5.319),*CCND1*(-1.599),*CDC25A*(-1.770),*CDCA7L*(-3.527),*CDKN1B*(+1.821),*CDKN2C*(-2.195),*CFL1*(-1.479),*E2F1*(-2.816),*E2F2*(-5.760),*ECT2*(-2.186),*FBXW7*(+1.413),*FOSL1*(-1.405),*HIPK3*(+1.979),*HMGB3*(-1.637),*HMGN2*(-4.630),*HMMR*(-1.998),*KIF14*(-2.677),*MKI67*(-6.492),*MLXIPL*(-1.570),*NCAPH*(-4.978),*NR4A3*(+3.464),*NUF2*(-2.814),*PCNA*(-1.843),*PFAS*(-1.680),*PTGS2*(+3.339),*PXMP2*(-2.217),*RACGAP1*(-4.835),*RASD1*(+2.025),*SDC1*(-2.179),*SERPINE1*(-1.537),*SKA3*(-3.640),*SPP1*(+2.156),*STIL*(-2.869),*TESMIN*(-2.077),*USP13*(-1.819),*VEGFA*(+1.835) |
|  | ligand-dependent nuclear receptor | PPARA | 9.37E-04 | 0.735 | 45 | 14 | 31 | *ABCA1*(+2.225),*ABCB4*(+2.122),*AURKA*(-4.407),*BCL2L1*(-1.951),*C3*(+32.006),*CCNA2*(-5.214),*CCNB1*(-5.319),*CCND1*(-1.599),*CCNE1*(-1.820),*CDK1*(-2.605),*CDKN2C*(-2.195),*CFH*(+1.599),*CHAF1B*(-4.714),*CYP51A1*(-1.402),*ECI1*(-1.439),*GLUD1*(-1.450),*H2AFX*(-3.444),*H2AFZ*(-1.900),*HADH*(-1.510),*HIST1H1C*(-3.279),*HLA-E*(+1.366),*HMGCS1*(+1.490),*KIF20A*(-7.937),*KIF2C*(-5.609),*KIF4A*(-5.155),*MAD2L1*(-2.803),*MKI67*(-6.492),*MSMO1*(+1.603),*NPPB*(-2.715),*OGG1*(-1.517),*PBLD*(+1.834),*PCNA*(-1.843),*PCTP*(+1.678),*PLK1*(-8.213),*PRC1*(-5.844),*PTGS2*(+3.339),*QPCT*(+1.656),*RTN4*(+1.438),*SC5D*(+2.024),*SMC2*(-1.640),*SMC4*(-1.901),*STIM1*(-1.452),*TOP2A*(-4.327),*VEGFA*(+1.835) |
|  | ligand-dependent nuclear receptor | PPARG | 1.64E-03 | -1.202 | 44 | 21 | 23 | *ABCA1*(+2.225),*APH1B*(+2.054),*ARL4D*(-1.503),*BCL2L1*(-1.951),*BCL6*(+1.817),*BDH1*(-1.539),*BECN1*(+1.368),*BIRC5*(-3.100),*BRCA1*(-2.980),*C3*(+32.006),*CCND1*(-1.599),*CCPG1*(+2.465),*CDKN2C*(-2.195),*CTPS1*(-1.576),*FGF1*(-2.606),*FOSL1*(-1.405),*IDH1*(+1.659),*LAPTM4A*(+1.631),*MCM7*(-4.228),*MCTP1*(+1.563),*MKI67*(-6.492),*MKNK2*(+1.716),*MMP14*(+1.527),*MRTO4*(-1.429),*NDUFA5*(+2.420),*NPPB*(-2.715),*OXR1*(+1.542),*PCNA*(-1.843),*PCTP*(+1.678),*PMM1*(-1.567),*PTGS2*(+3.339),*RGL1*(+1.418),*SDC1*(-2.179),*SERPINE1*(-1.537),*SPP1*(+2.156),*STAT5B*(-1.637),*STIM1*(-1.452),*TAGLN*(-2.352),*TGFBR1*(+1.595),*TGFBR2*(+2.235),*TSC22D3*(+1.632),*TUBB*(-1.883),*VEGFA*(+1.835),*­*(-1.608) |
| S6K1sh | other | TXNIP | 1.67E-02 | 1.273 | 5 | 2 | 3 | CCNA2(-1.396),DDIT4(+1.814),FASN(-1.840),LPL(-1.488),PTGS2(+1.872) |
|  | kinase | PERK | 2.58E-02 | 1.154 | 8 | 6 | 2 | ATF5(-1.426),BIRC3(+1.777),FBXO8(+2.144),HERPUD1(+1.435),LMO4(+1.479),PPP1R15A(+1.460),PRDM1(+2.026),TP53(-1.515) |
|  | transcription regulator | ATF4 | 5.67E-03 | 1.532 | 12 | 7 | 5 | APBA3(+1.898),ATF5(-1.426),CHAC1(+2.133),DDIT4(+1.814),FASN(-1.840),HERPUD1(+1.435),PMAIP1(+1.878),PPP1R15A(+1.460),PRKDC(-1.401),PTGS2(+1.872),PYCR1(-1.283),RBPJ(-1.709) |
|  | chemical drug | troglitazone | 4.48E-04 | -0.823 | 26 | 7 | 19 | ACADS(-1.516),ANGPTL4(+2.606),AR(-2.507),BIRC3(+1.777),BIRC5(-1.550),CCNB1(-2.235),CDCA7L(-1.552),FASN(-1.840),HIPK2(-1.735),HIPK3(+1.479),HMGN2(-1.913),IRS1(-1.440),LPL(-1.488),MKI67(-1.460),NCAPH(-2.325),NNMT(-1.512),NRIP1(+1.387),PTGS2(+1.872),RACGAP1(-1.726),SDC1(-1.454),SERPINE1(+1.270),SKP2(-3.178),SLC27A1(-1.698),TBXAS1(-1.494),TP53(-1.515),WEE1(+1.396) |
|  | ligand-dependent nuclear receptor | PPARA | 7.43E-04 | -1.428 | 29 | 3 | 26 | ACADS(-1.516),ANGPTL4(+2.606),AURKA(-1.669),CCNA2(-1.396),CCNB1(-2.235),CHAF1B(-1.859),CS(-1.430),FADS1(-1.473),FASN(-1.840),FBXO21(-1.563),GOT2(-1.368),GPD2(-1.559),H2AFX(-1.446),HADH(-1.502),HIST1H1C(-1.519),IL10(+2.121),KIF20A(-2.078),KIF2C(-1.979),KIF4A(-1.452),KYAT1(-2.087),LPL(-1.488),MKI67(-1.460),PLK1(-2.115),PRC1(-1.731),PTGS2(+1.872),SLC27A1(-1.698),SORD(-1.721),SRM(-1.801),TOP2A(-1.614) |
|  | ligand-dependent nuclear receptor | PPARG | 5.03E-05 | 0.512 | 33 | 13 | 20 | ACADS(-1.516),ADRB2(-1.714),ANGPTL4(+2.606),BACE1(-1.485),BIRC5(-1.550),CDK6(+1.806),CS(-1.430),ESRRA(+1.343),FASN(-1.840),HEBP1(-1.634),HIVEP2(+1.317),IL10(+2.121),IRS1(-1.440),KLF6(+1.330),LHPP(-1.995),LPL(-1.488),MCM7(-1.633),MKI67(-1.460),PRDM1(+2.026),PTGS2(+1.872),RHOB(+1.494),RPSA(-1.302),SDC1(-1.454),SERPINE1(+1.270),SLC27A1(-1.698),SLC44A1(-1.829),TAGLN(+1.682),TBXAS1(-1.494),TKT(-1.370),TP53(-1.515),VAMP2(+1.562),VDR(+2.193),WIPF1(-1.448) |
|  | ligand-dependent nuclear receptor | PPARD | 5.65E-02 | -2.263 | 12 | 3 | 9 | ANGPTL4(+2.606),BIRC5(-1.550),FASN(-1.840),GOT2(-1.368),GPD2(-1.559),IL10(+2.121),KYAT1(-2.087),LPL(-1.488),PTGS2(+1.872),SLC27A1(-1.698),SORD(-1.721),TP53(-1.515) |
|  | complex | PPARÎ±-RXRÎ± | 1.25E-02 |  | 3 | 0 | 3 | GOT2(-1.368),LPL(-1.488),SLC27A1(-1.698) |

RNA sequencing data were analyzed by IPA.

Type: regulator type

Regulator: targets of this regulator are overrepresented in the list

*P*: nominal p-value of the overrepresentation

Z: z-score of the prediction – positive for activated, negative for inhibited function.

N: number of regulator’s target genes in the list

pos: number of upregulated molecules

neg: numbets of downregulated molecules

Molecules: genes (or complexes) from the gene list known to be regulated
