## Supplementary Table 6 for "Selective abrogation of S6K2 maps lipid homeostasis as a survival vulnerability in MAPKi-resistant NRAS^MUT^ melanoma"

**Table S6. qRT-PCR primer sequences**

| Primer: ACOX1 Forward:  AAT CGG GAC CCA TAA GCC TTT | Primer: ACOX1 Reverse: GGG AAT ACG ATG GTT GTC CAT TT |
| --- | --- |
| Primer: ACOX3 Forward:  TCG CTC CTC CTG ACT TTG TT | Primer: ACOX3 Reverse:  CCA GTC CCA CTA GAG CTT CG |
| Primer: ACSL1 Forward:  CTG CCC CAG ATC AGT TCA TT | Primer: ACSL1 Reverse:  GCC TTC TCT GGC TTG TCA AC |
| Primer: ACSL4 Forward:  CCC CTG AAA CTG GTT TGG TA | Primer: ACSL4 Reverse:  AAA CCA CCA GGC TAC CTC CT |
| Primer: ACTB Forward:  AGC ACT GTG TTG GCG TAC AG | Primer: ACTB Reverse:  AGA GCT ACG AGC TGC CTG AC |
| Primer: CPT1A Forward:  CAA GGA CAT GGG CAA GTT TT | Primer: CPT1A Reverse:  ATG CTT CTC AGA CGC CAA CT |
| Primer: PGC1α Forward:  GAGTCTGTATGGAGTGACATCG | Primer: PGC1α Reverse:  TCACTGCACCACTTGAGTCC |
| Primer: PPARα Forward:  CTA TCA TTT GCT GTG GAG ATC G | Primer: PPARα Reverse:  AAG ATA TCG TCC GGG TGG TT |
| Primer: PPARß Forward:  GTC ACA CAA CGC TAT CCG TTT | Primer: PPARß Reverse:  AGG CAT TGT AGA TGT GCT TGG |
| Primer: PPARγ Forward:  CGT GGC CGC AGA TTT GAA | Primer: PPARγ Reverse:  CTT CCA TTA CGG AGA GAT CCA C |
| Primer: S6K1 Forward:  ACA TAG ACC TGG ACC AGC CA | Primer: S6K1 Reverse:  CTC ACA ATG TTC CAT GCC AAG TT |
| Primer: S6K2 Forward:  GGA TTT GGA GAC GGA GGA AGG | Primer: S6K2 Reverse:  TTC ACG CTG GTC TCA GTC AG |
| Primer: XBP-1 (spliced) Forward:  TGC TGA GTC CGC AGC AGG TG | Primer: XBP-1 (spliced) Reverse:  GCT GGC AGG CTC TGG GGA AG |
